# Population histories of the United States revealed through fine-scale migration and haplotype analysis

**DOI:** 10.1101/577411

**Authors:** Chengzhen L. Dai, Mohammad M. Vazifeh, Chen-Hsiang Yeang, Remi Tachet, R. Spencer Wells, Miguel G. Vilar, Mark J. Daly, Carlo Ratti, Alicia R. Martin

## Abstract

The population of the United States is shaped by centuries of migration, isolation, growth, and admixture between ancestors of global origins. Here, we assemble a comprehensive view of recent population history by studying the ancestry and population structure of over 32,000 individuals in the US using genetic, ancestral birth origin, and geographic data from the National Geographic Genographic Project. We identify migration routes and barriers that reflect historical demographic events. We also uncover the spatial patterns of relatedness in subpopulations through the combination of haplotype clustering, ancestral birth origin analysis, and local ancestry inference. Examples of these patterns include substantial substructure and heterogeneity in Hispanics/Latinos, isolation-by-distance in African Americans, elevated levels of relatedness and homozygosity in Asian immigrants, and fine-scale structure in European descents. Taken together, our results provide detailed insights into the genetic structure and demographic history of the diverse US population.

## Introduction

The United States population is a diverse collection of global ancestries shaped by migration from distant continents and admixture of recent migrants and Native Americans. Throughout the past few centuries, continuous migration and gene flow have played major roles in shaping the diversity of the US. Mixing between groups that have historically been genetically and spatially distinct have resulted in individuals with complex ancestries, while within-country migration have led to genetic differentiation.^1–9^

Deeply characterizing population history is important for understanding human evolution and demographic history, as well as for adequate study design when associating genotypes to phenotypes.^10–13^ Earlier population genetic studies in the US broadly characterized this structure typically using a limited set of ancestry-informative markers or uniparental mtDNA and Y chromosome DNA data.^14^ As the cost of genetic technologies have dropped, more recent studies have inferred population history with more complete genome-wide data, typically using more than 100,000 SNPs ascertained via sequencing or genotyping.

Previous genetic studies of the US population have sought to infer genetic ancestry and population history primarily in European Americans, African Americans, and Hispanics/Latinos.^7–9, 15, 16^ European American ancestry is characterized by substantial mixing between different ancestral European populations and, to a lesser extent, admixture with non-European populations.^6, 8, 9^ Isolation among certain European population, such as Ashkenazi Jewish, French Canadian, and Finnish populations, have also resulted in founder effects.^17–19^ The mixing of European settlers with Native Americans have contributed to large variations in the admixture proportions of different Hispanic/Latino populations.^1, 4, 9^ Among Hispanics/Latinos, Mexicans and Central Americans have more Native American ancestry; Puerto Ricans and Dominicans have more African ancestry; and Cubans have more European ancestry.^1, 4^ In African Americans, proportions of African, European, and Native American ancestry vary across the country and reflect migration routes, slavery, and patterns of segregation between states.^2, 3, 7, 9, 20^ Although much effort has been made to understand the genetic diversity in the US, fine-scale patterns of demography, migration, isolation, and founder effects are still being uncovered with the growing scale of genetic data, particularly for Latin American and African descendants with complex admixture histories.^21, 22^ At the same time, there has been little research on the population structure of individuals with East Asian, South Asian, and Middle Eastern ancestry in the US.

Many previous studies have investigated specific population histories in the US at relatively small scales–on the order of hundreds to thousands of individuals. These studies have provided deep insights into many specific populations, with some well-powered to infer population history across a breadth of ancestries. Some of these insights have been made by applying methods which are only computationally tractable at smaller scales.^23, 24^ More recently, however, important insights highlight the need for broader and more comprehensive investigations of population history. For example, recent studies have shown that population structure is inaccurately captured in small sample sizes.^10, 13^ Additionally, millions of Americans have been interested enough in their genetic ancestry to pay direct-to-consumer companies for individual-level genetic ancestry reports.^8, 9^ The reliability of these reports is high for many individuals, but they are dependent on 1) the representativeness of their reference panel or customer database, 2) completeness and accuracy of multigenerational birth origin data, and 3) the application of multiple approaches to gain holistic insights into population history.

In this study, we comprehensively evaluate the population history of over 32,000 genotyped individuals in the US who partook in the National Geographic Genographic Project, a not-for-profit public participation research initiative to study human migration history.^25^ This project has several distinct advantages compared to other large-scale population genetics datasets. Individual-level genetic data are accessible to researchers around the world to answer anthropological questions. Additionally, most participants report birthplace and ethnicity data for themselves, their parents, and their grandparents, enabling fine-scale insights into recent history. Furthermore, participants report their postal code when they participated in the study, enabling analysis of intragenerational migration. These data therefore enable high spatiotemporal resolution into historical migration patterns. While these trends are consistent with US history at the population scale, we note that genetic ancestry patterns are not commensurate with individual-level ethnicity (i.e. cultural identity).

Here, we leverage these advantages over existing data to identify patterns of genetic ancestry by studying pairwise sharing among the project participants. We combine these comparative patterns with ancestral birth origin records and geographic information to uncover recent demographic and migration trends. By comprehensively analyzing these data to learn about recent migration events, we gain deeper insights into ancestral origins than in many existing studies, especially into Latin America. We also provide early insights into Asian Americans often ignored in genetic studies of the US, including South Asians, East Asians, and Middle Easterners. We also identify detailed patterns among European and African American populations, recapitulating some similar trends reported previously. Taken together, we use accessible individual-level genetic and birth record data to provide insights into the ancestral origins and complex population histories in the US.

## Results

### Genetic ancestry and diversity across the United States

To assess the diversity of ancestries among individuals in the Genographic Project, we first performed principal component analysis, projecting Genographic samples into the same principal component (PC) space as that of the 1000 Genome Project samples (**Figure 1A-C; Figure S1-S2**).^26, 27^ Since self-reported ancestry was not consistently provided across all Genographic individuals, we leveraged the 1000 Genomes Project data to assign continental ancestry to each Genographic sample (**Methods; Supplemental Materials and Methods**). We first trained a Random Forest classifier on the first 10 principal components (PCs) of the 1000 Genome Project samples with super population groupings as ancestry labels (EUR = European, AMR = Admixed American, AFR = African, EAS = East Asian, SAS = South Asian). We then used the trained model to assigned continental ancestry to each individual in the Genographic cohort at >90% confidence. A total of 3,028 individuals (9.3% of total) did not meet the classification threshold (**Figure 1C; Table S1**). The inability to classify these individuals may be due to the complex and variable admixture profiles of certain populations such as Hispanics/Latinos.

**Figure 1.**
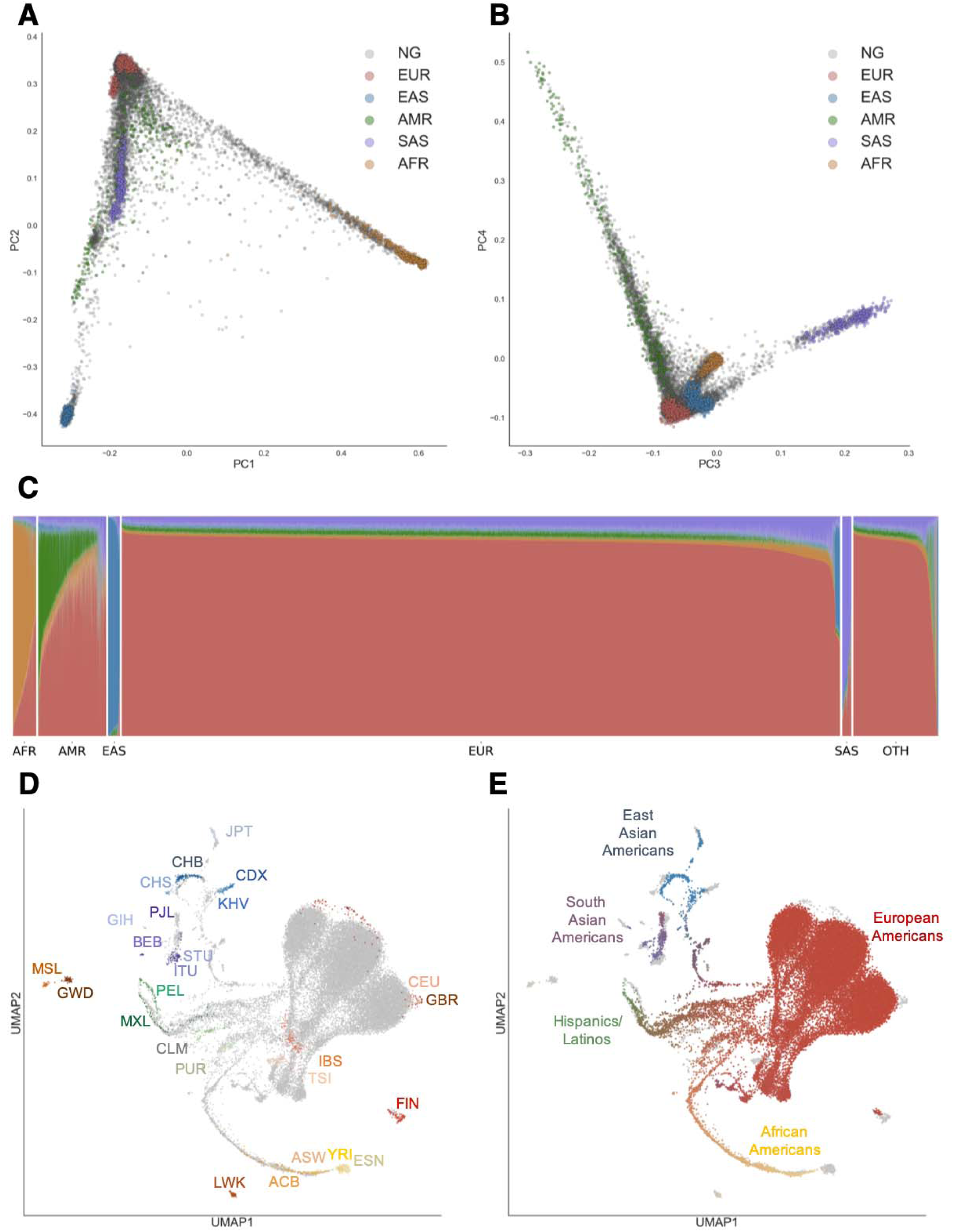
Genetic Diversity of the US Population. (A) Principal Components Analysis (PCA) of individuals in the United States and in the 1000 Genome Project. Each individual is represented by a single dot. Individuals in this study are colored in grey while 1000 Genome Population individuals are colored by super population (EUR = red, AFR = yellow, NAM = green, EAS = blue, SAS = purple). Principal components (PC) 1 and PC 2 are shown. (B) Similar to (A), except principal components (PC) 3 and PC 4 are shown. (C) ADMIXTURE analysis at K=5 of individuals in this study. Each individual was assigned a continent-level ancestry label using a Random Forest model trained on the super population labels and the first 10 PCs of the 1000 Genome Project dataset. OTH = individuals who did not meet the 90% confidence threshold for classification. (D) – (E) UMAP projection of the first 20 PCs. Each dot represents one individual. In (D), individuals in the 1000 Genomes Project are colored by population while US individuals are in grey. In (E), 1000 Genome Project individuals are colored in grey while US individuals are colored based on their admixture proportions from ADMIXTURE. The color for each dot was calculated as a linear combination of each individual’s admixture proportion and the RGB values for the colors assigned to each continental ancestry (EUR = red, AFR = yellow, NAM = green, EAS = blue, SAS = purple). See Materials and Methods for specific population labels.

Regional differences in genetic ancestry correspond to historical demographic trends. We evaluated the distributions of classified individuals across the four designated US Census regions: South, Northeast, Midwest, and West (**Table S1**). Classified individuals of European descent make up the majority (78.5%) of the Genographic cohort and are the most prevalent in the Midwest (82.8% of individuals in the Midwest; P<0.01, Fisher’s exact test; **Table S1**). Individuals of Native American ancestry are most prominent in the West and South (9.7% and 7.8% of total individuals in the West and South, respectively; P<0.05, Fisher’s exact test). Individuals classified as having African ancestry are most common in the South (3.2%), followed by the Northeast (3.0%). East Asians mostly reside in the West (2.1%), while South Asians are most abundant in the Northeast (1.0%). While the proportion of individuals classified as of European descent in the Genographic cohort (78.5%) are similar to the proportions of individuals reported as “White” in the U.S. Census Data (76.1%; **Table S2**), we note that genetic ancestry is not a direct measure of ethnicity and race, and the two are not fully comparable (**Supplemental Materials and Methods**). The large proportion of unclassified individuals also hinders our ability to properly compare the Genographic cohort to the US census and understand how representative the Genographic cohort is of the US population.

To uncover population substructure, we performed dimensionality reduction with Uniform Manifold Approximation and Projection (UMAP) on the first 20 PCs of a combined Genographic and 1000 Genomes Project dataset.^28, 29^ By leveraging multiple PCs at once, UMAP can disentangle subcontinental structure (**Figure 1D-E; Figure S3-S4**). Similar to previous analysis,^29^ populations in the 1000 Genomes Project form distinct clusters corresponding to ancestry and geography. The Genographic individuals project into several clusters, overlapping with the 1000 Genomes Project clusters. Consistent with the PCA and ADMIXTURE analysis, the largest clusters correspond to European ancestry and cluster closely with the 1000 Genomes CEU and GBR populations (CEU=Utah Residents with Northern and Western European Ancestry, GBR=British in England and Scotland).

While UMAP is a visualization tool with no direct interpretation of genetic distance, the continuum of points connecting UMAP clusters reflects the varying degrees of estimated admixture between different continental ancestries. In particular, the complex population structure of Hispanics/Latinos is shown by the points spanning between the clusters of European, Native American, and African ancestry. Coloring of these points based on ancestry proportions affirms the relationship between the degree of admixture and their relative position between reference clusters. Interestingly, African American individuals from both datasets form a single continuum from the European cluster to the Yoruba (YRI) and Esan (ESN) populations of Nigeria in the 1000 Genomes Project, indicative of the West African origins of most African Americans. This observation is consistent with and further expands the previous finding that the African tracts in the admixed 1000 Genomes populations of ACB and ASW were previously found to be similar to the Nigerian YRI and ESN populations.^2, 30^

### Fine-scale structure among US individuals of Asian ancestry

Existing genetic studies of the US population have largely overlooked East Asian and South Asian populations, likely due to their underrepresentation in datasets. We therefore explored the population structure of Genographic Project individuals classified as East Asians and South Asians. We applied fineSTRUCTURE to hierarchically cluster unrelated individuals in each population based on patterns of shared ancestry and inferred a total of 40 East Asian clusters (**Figure 2A**) and 26 South Asian clusters (**Figure 2B**). These clusters further organized into clades on the tree to reveal broader genetic structure. To further visualize these structures, we performed PCA on the fineSTRUCTURE coancestry matrix. Compared to traditional PCA, distinctions between groups of individuals were clearer with fineSTRUCTURE PCA, particularly at the broader levels of genetic differentiation (**Figure 2A** and **S5A**; **Figure 2B** and **S6A;** Methods). We also estimated subcontinental admixture proportions with ADMIXTURE using the East Asian and South Asian populations in the 1000 Genomes Project and the Human Genome Diversity Project (HGDP) as reference populations (**Figure S5B-C, S6B-C**). Finally, we leveraged data from individuals who provided grandparental birth origin to help annotate and interpret the clusters and clades.

**Figure 2.**
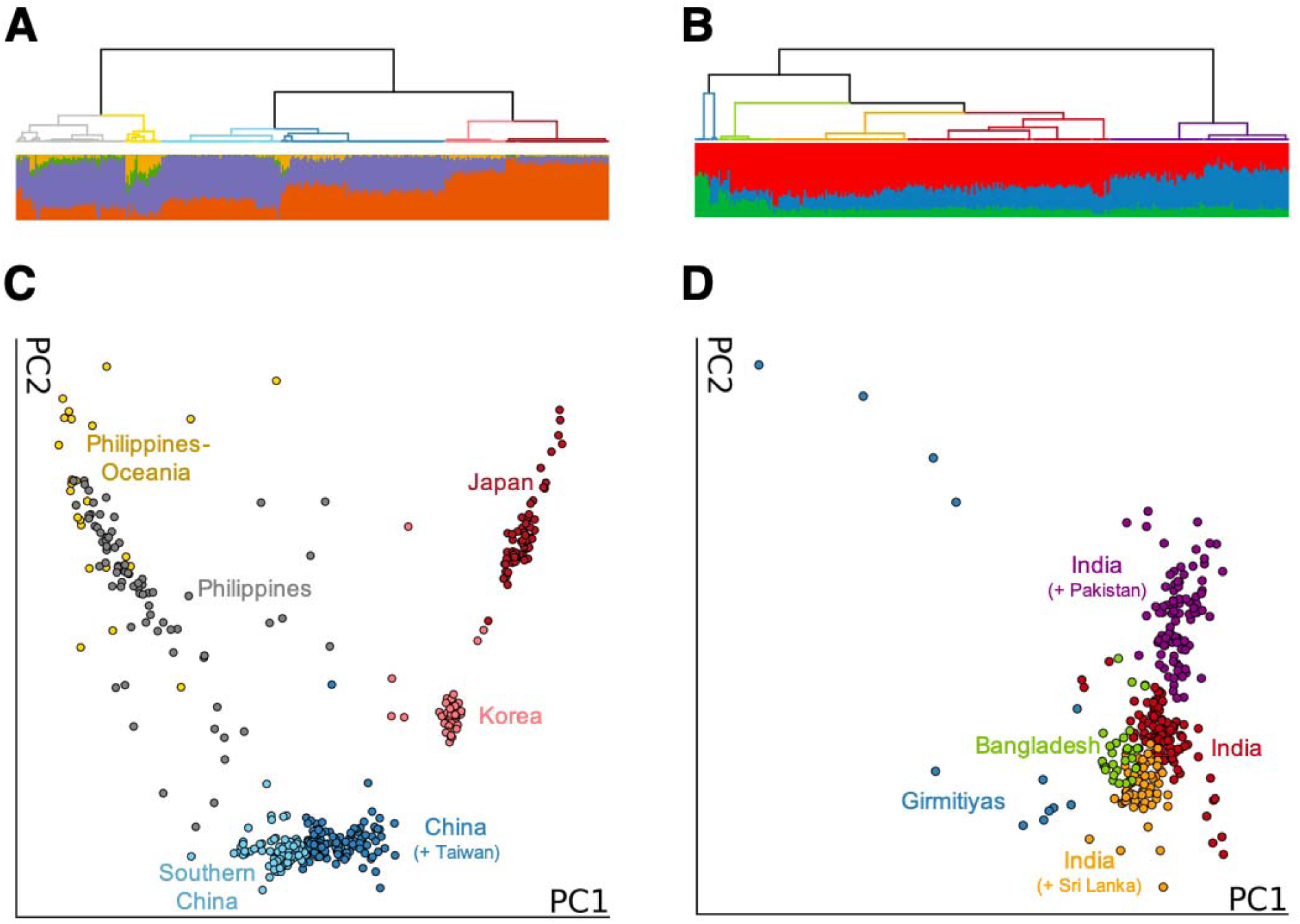
Population Structure of East Asian and South Asian Individuals in the US. (A) – (B) fineSTRUCTURE dendrogram showing the hierarchical relationship between clusters inferred using the genotypes of classified East Asian individuals (A) and South Asian individuals (B). Branch colors represent clades with shared ancestral origins. The admixture proportion of each individual is displayed as a bar plot in the corresponding position below the dendrogram. The number of ancestral populations, K, is four for East Asians (A) and three for South Asians (B). (C) – (D) Principal component analysis (PCA) of the fineSTRUCTURE co-ancestry matrix. Each individual (point) corresponds to a Genographic individual classified as either East Asian (C) or South Asian (D). The color of each point corresponds to a clade in the fineSTRUCTURE dendrogram shown in (A) and (B).

The patterns of shared ancestry among these US individuals capture the genetic diversity of East Asia and South Asia. The East Asian clusters broadly organize into six major clades, reflecting the different countries of ancestral origin (**Figure 2A**). At the highest level of genetic differentiation (top level of the hierarchical tree), individuals from Southeast Asia separate from East Asians. This Southeast Asian clade is predominantly represented by Filipinos with a branch of individuals with more Oceanic origins (shown in grey and yellow, respectively). Admixture proportions vary among the Southeast Asian individuals, likely due to the large number of ethnolinguistic groups that are found in the Philippines and neighboring islands. The East Asian clade further separates into individuals of Chinese descent (light blue and dark blue) and those from Japan (dark red) and Korea (light red). While the two Chinese-related groups share a branch on the tree, Taiwanese ancestral origins are more prevalent in one of the groups (dark blue), the ‘China (+ Taiwan)’ group, while the other group (light blue), labeled ‘Southern China’, also contains some individuals from Laos and Vietnam. Lower levels of hierarchy did not differentiate these ancestral origins into separate groups. PCA and ADMIXTURE analysis for these two groups show that the ‘China (+ Taiwan)’ cluster resembles the Han Chinese (CHB) population in the 1000 Genome Project while the ‘Southern China’ group resembles the Southern Han Chinese (CHS) population (**Figure S5**). Among the South Asian individuals, we observed genetic differentiation between individuals with ancestral origins from India, reflecting the diverse population structure previously observed in India.^1, 31^ Of the three clades with majority Indian ancestral origin, ancestral origins from Pakistan was observed in the ‘India (+ Pakistan)’ clade, while Sri Lankan ancestral origins were present in the ‘India (+ Sri Lanka)’ clade. Individuals in these two clades resemble the Punjabi from Lahore, Pakistan (PJL) and Sri Lankan Tamil (STU) populations in the 1000 Genomes Project, respectively (**Figure S6**). Similarly, we also find a clade of individuals with Bangladesh ancestral origins that is similar to the 1000 Genomes Project Bengali from Bangladesh (BEB). Interestingly, we also inferred a small, but genetically distinct ‘Girmitiyas’ clade (N = 12; blue branch in **Figure 2B**). While the small sample size makes it difficult to accurately assess this clade, we note that many former British colonies (e.g. Trinidad and Tobago, Fiji, Barbados, Guyana) are represented in the ancestral origins of these individuals. We therefore hypothesize that these individuals may potentially be descendants of Girmitiyas, indentured Indian laborers brought to those former colonies.^32^

### Population differentiation and migration rate inference across the United States

Understanding the relationship between genetics and geography can provide insights into demographic history. Previous analyses of this relationship in the US population have primarily compared data aggregated at the state or regional level.^7, 9^ Such approaches, however, do not capture the fine-scale patterns of genetic similarity that are not influenced by discrete administrative boundaries. We therefore sought to infer continuous population structure across space with the estimating effective migration surfaces (EEMS) method.^33^ EEMS statistically measures effective migration rates by overlaying a dense grid of evenly-spaced demes and calculating genetic differentiation (i.e. resistance distance) between neighboring demes. Higher rates of migration are inferred in locations where genetic similarity is high (colored in blue) while lower rates of migration are inferred in locations where genetic similarity is low (these locations are also referred to as migration barriers and colored in dark orange). We applied EEMS to genetically classified Europeans, African Americans, and Hispanic/Latinos across the contiguous 48 states. We excluded East Asians and South Asians due to low sample density.

The inferred migration rates for African Americans reveal genetic signatures of historical demographic events (**Figure 3A; Figure S7**). Along the Atlantic coast from the Florida Panhandle to southern Maine, genetic similarity and effective migration rates are relatively high, indicating the constant migration and similar effective population sizes of African Americans in these states. However, we also observe a strong north-south barrier to migration starting along the Appalachian Mountain Range, continuing north up the Mississippi River, and extending west across the rest of the country. This migration barrier, along with the migration barrier spanning Texas and New Mexico, reveals a pattern of genetic relatedness across geography that is consistent with the Great Migration from the 1910s to the 1960s in which an estimated 6 million African Americans migrated out of the South to cities across the Northeast, Midwest and West.^7, 34^

**Figure 3.**
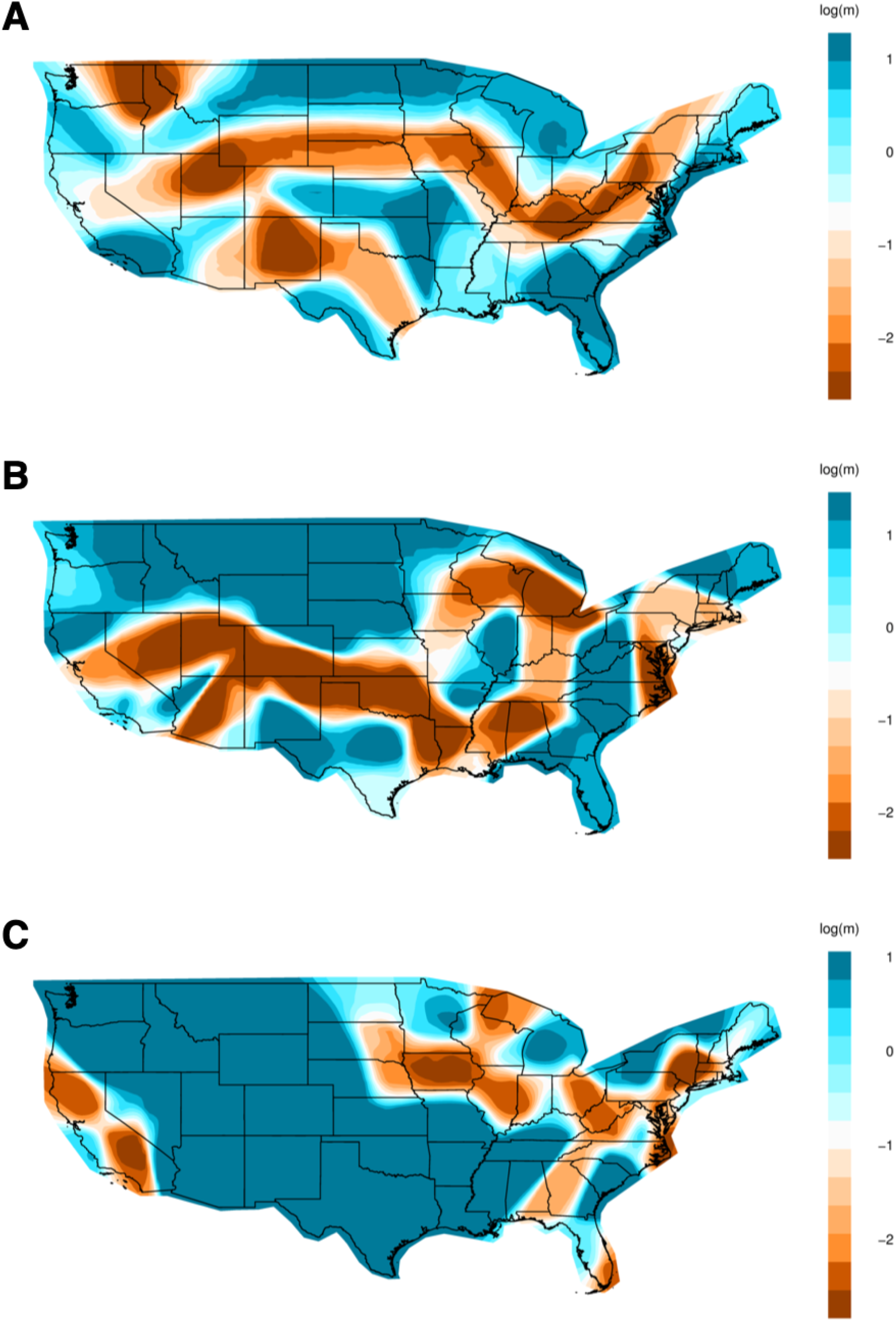
Migration Rates of African Americans, Hispanics/Latinos, and Europeans within the United States. (A) – (C) Migration rates inferred with EEMS for African Americans (A), Hispanics/Latinos (B), and Europeans (C). EEMS models the relationship between genetics and geography by assessing the decay of genetic similarity with respect to geographic distance. Colors and values correspond to inferred rates, *m*, relative to the overall migration rate across the country. Shades of blue indicate higher migration (i.e. log(m) = 1 represents effective migration that is tenfold faster than the average) and higher levels of genetic similarity while shades of orange indicate migration barriers and lower levels of genetic similarity.

A highly complex pattern of genetic similarity exists amongst present-day Hispanics/Latinos across the country, capturing regional genetic structure. Across the southwestern states, two regions bordering Mexico—one in California and another extending from New Mexico to Texas—exhibit high levels of genetic similarity and effective migration rates (**Figure 3B; Figure S7**). Separated by a migration barrier in Arizona, these two distinct regions likely reflect known differences in the northward migration from east versus west Mexico.^8, 35^ High genetic similarity and relative rates of effective migration are also observed in Florida and continue northward. However, barriers to migration are observed in states immediately east of the Mississippi River, the likely resulting from varying degrees of admixture.

The patterns of genetic similarity for Europeans capture subcontinental structure. With the exception of the states in the Midwest and along the Atlantic coast, elevated levels of genetic similarity and relative migration rates are observed across most of the country. We find low effective migration rates surrounding Minnesota and Michigan, likely due to the genetic dissimilarity of Finnish and Scandinavian ancestry that is abundant in the region (**Figure 3C; Figure S7**).^8^ We also find reduced migration rates across Ohio, West Virginia, and Virginia, suggesting the existence of genetic differentiation along the Appalachian Mountains. Many of the major cities, such as Washington, DC, Philadelphia, and Miami, also exhibit low genetic similarity, perhaps due to greater genetic diversity and admixture within cities. The migration barrier and lower genetic similarity encompassing metropolitan New York City may be explained in part by the large presence of divergent European populations, such as Ashkenazi Jews, in those areas (**Figure 3C**).

### Coupling fine-scale haplotype clusters and multigenerational birth records uncovers distinct subcontinental structure

To disentangle more recent and subtle population structure, we performed identity-by-descent (IBD) clustering on the Genographic cohort and annotated clusters using multigenerational self-reported birth origin data. We first built an IBD network from pairwise IBD sharing among 31,783 unrelated individuals, where vertices represent individuals and edges represent the cumulative IBD (in centimorgans, cM) between pairs of individuals. We employed the Louvain method, a greedy heuristic algorithm, to recursively partition vertices in the graph into clusters that maximize modularity for each iteration.^8, 36^ The clusters of individuals resulting from each iteration can be interpreted as having greater amounts of cumulative IBD shared between individuals within the cluster than with those outside of the cluster. To aid in the interpretation of the clusters, we merged clusters with low genetic differentiation (F_ST_ < 0.0001), resulting in a final set of 25 clusters (**Table 1**). We annotated each cluster based on ancestral birth origin and ethnicity data and constructed a neighbor-joining tree based on the F_ST_ values (**Figure S8)**. 98% of the 3,028 individuals that were not classified by our Random Forest model were assigned to a haplotype cluster. No single cluster was overrepresented by unclassified individuals, as unclassified individuals comprised of 8-11% of each cluster.

**Table 1.** Summary of Haplotype Clusters. Cumulative runs of homozygosity (cROH) was calculated by summing the regions of continuous homozygous segments. Cumulative IBD was determined by summing IBD segments of ≥ 3 cM and filtering for only pairs ≥ 12cM and ≤ 72 cM. Statistics were determined within haplotype clusters, rather than across the ancestrally heterogeneous and imbalanced full network.

| **Cluster** | **Samples**<br>**(count)** | **Median**<br>**Cumulative ROH**<br>**(cM)** | **Median**<br>**Cumulative IBD**<br>**(cM)** |
| --- | --- | --- | --- |
| Northwest Europe 1 | 11,725 | 2.88 | 15.23 |
| Northwest Europe 2 | 1,571 | 2.80 | 15.15 |
| Ireland | 2,137 | 2.85 | 15.42 |
| Central Europe | 3,116 | 2.83 | 15.06 |
| Eastern Europe | 2,471 | 3.16 | 15.37 |
| Southern Europe | 1,626 | 2.73 | 14.98 |
| Italy | 697 | 6.91 | 14.64 |
| Greece-Italy | 238 | 7.28 | 15.02 |
| Scandinavia | 717 | 3.02 | 15.54 |
| Finland | 314 | 3.67 | 17.50 |
| Acadia | 249 | 3.89 | 19.48 |
| French Canadian | 314 | 2.89 | 16.60 |
| Ashkenazi Jewish | 1,475 | 11.26 | 31.75 |
| Admixed Jewish | 445 | 2.75 | 15.50 |
| Hispanics/Latinos | 810 | 3.53 | 16.38 |
| Hispanics/Latinos in California | 573 | 4.10 | 17.11 |
| Hispanics/Latinos in New Mexico | 163 | 5.52 | 21.92 |
| Hispanics/Latinos in Texas | 177 | 6.27 | 23.65 |
| Puerto Rico | 350 | 8.01 | 26.23 |
| African Americans South | 761 | 3.34 | 19.56 |
| African Americans North | 420 | 2.94 | 15.90 |
| East Asia | 561 | 3.65 | 19.63 |
| Southeast Asia | 325 | 8.44 | 17.90 |
| South Asia | 389 | 10.42 | 14.82 |
| Greater Middle East | 93 | 9.01 | 17.16 |

Genetic and geographic diversity is greatest amongst Hispanic/Latino haplotype clusters. We identified a total of five Hispanic-related clusters. The largest of these cluster (n=810) is strongly associated with south Florida (OR = 10.4; p = 2.5e-25; **Figure 4**, **Table S3**) but is also found in California, and Texas (OR ≥ 2; p < 0.05). No single ancestral birthplace characterizes this cluster, as the US, Mexico, and Cuba each make up more than 10% of the birth origin labels. Proportions of European ancestry tracts inferred with RFMix^37^ are higher in this cluster (mean = 72.7%, sd=20.4%) than in the other Hispanic/Latino clusters (mean = 48.0% – 67.4%). Puerto Ricans characterize a substantial proportion of another Hispanic/Latino cluster associated with Florida (OR > 4), as well as New York City (OR > 5). Unlike the other Hispanic clusters, the Puerto Rican cluster shares the same branch on the F_ST_ tree as the African American clusters (**Figure S8**), likely due to relatively high proportions of African ancestry (mean = 11.2%, sd = 9.0%) among Puerto Ricans.

**Figure 4.**
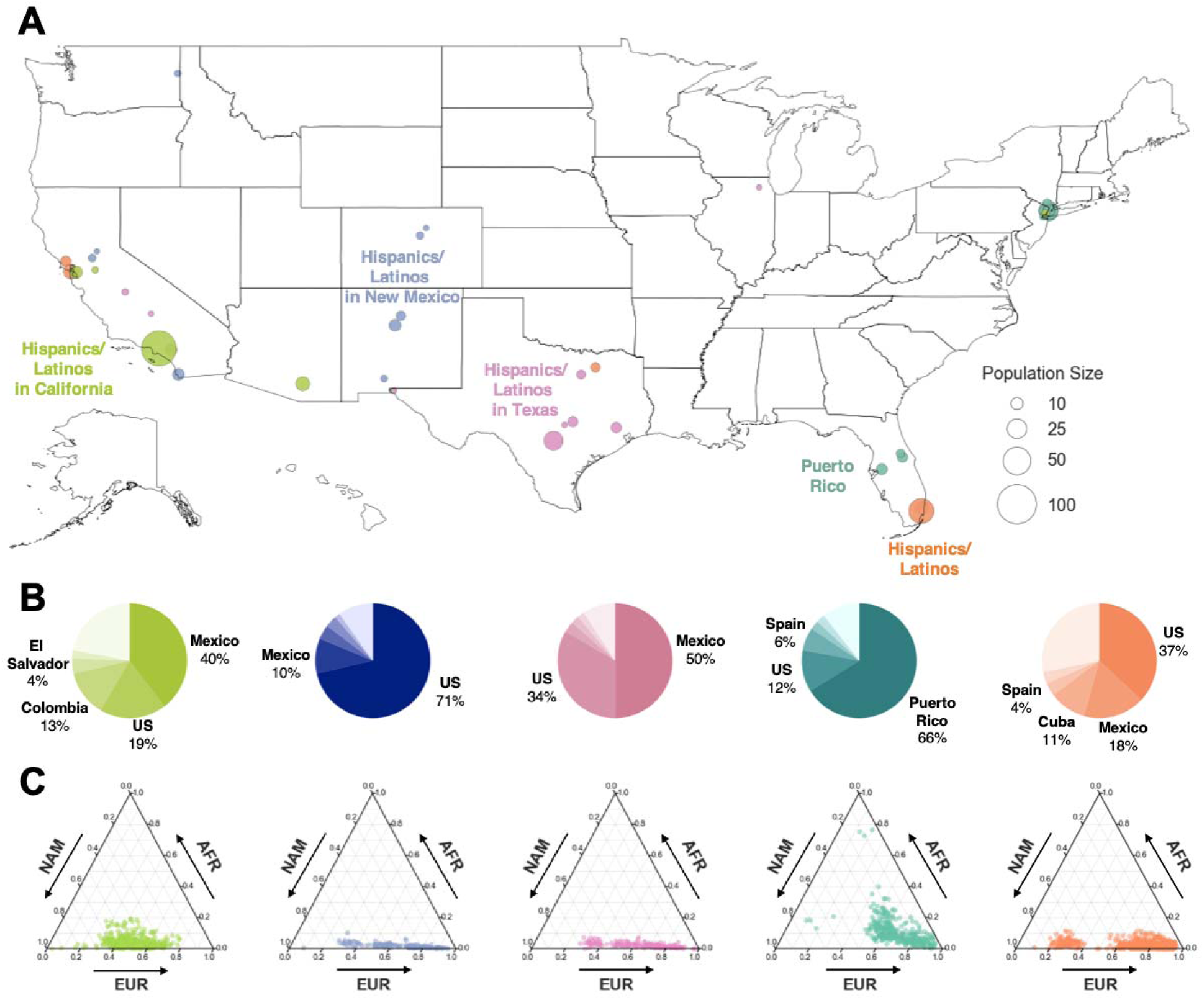
Distribution of Hispanic/Latino Haplotype Clusters. (A) Geographical distribution of Hispanic/Latino haplotype clusters. Each dot corresponds to a county containing present-day individuals and the size of the dot signifies the number of samples of the particular cluster in that county. Only the Hispanic/Latino cluster with the highest odds ratio is shown for each county, and for clarity, only the top ten locations with the highest odds ratio are shown for each cluster. Maps showing the full distribution for each haplotype cluster can be found in the supplement (Figure S11). (B) Ancestral birth origin proportions of each cluster for individuals with complete pedigree annotations, up to grandparent level. Proportions were calculated from aggregating the birth locations of all grandparents corresponding to members of each haplotype cluster. For each chart, only the top five birth origins are shown as individual proportions; the remaining birth origins are aggregated into one slice (lightest color). (C) Ternary plots of ancestry proportions based on local ancestry inference for each haplotype cluster. Each dot represents one individual.

Three distinct clusters of Hispanics were found in the Southwest (**Figure 4A**): one strongly associated with New Mexico (OR > 4; p < 0.05), another primarily in Texas (OR > 3; p < 0.05), and the third associated with Southern California (OR > 2; p < 0.05). Combined with the EEMS analysis, these clusters confirm our observation of parallel migration routes from east and west Mexico into Southwestern United States. While the genetic differentiation of these three clusters are subtle (F_ST_=0.001-0.003), ancestral birth origin patterns and local ancestry proportions for these clusters reveal meaningful dissimilarities. Whereas the majority of Hispanics in New Mexico report US ancestral origins, the recent ancestors of Hispanics in Texas are predominantly from Mexico. Nonetheless, these two clusters share similar local ancestry proportions with only slight genetic dissimilarity that result in a moderate decrease in migration rate (from darker blue to light blue in **Figure 2B**). Unlike the Hispanic clusters associated with New Mexico and Texas, the Hispanics in California cluster contain greater proportions of ancestors from Central and South American (e.g., Colombia and El Salvador). Proportions of Native American ancestry is also highest in this cluster (**Figure 4B**). Taken together, these two differences further explain the presence of the migration barrier in Arizona between the Hispanics in the California and the Hispanics in New Mexico.

Historical immigration of Europeans into the US occurred in successive waves, with Northern and Western Europeans making up one wave from the 1840s to 1880s and another wave comprising of Southern and Eastern Europeans occurring from the 1880s to 1910s.^38^ Consistent with this immigration pattern, haplotype clusters with ancestries from Northwest and Central Europe have higher proportions of US ancestral birth origins than haplotype clusters from Southern and Eastern Europe, suggesting earlier immigration (**Figure 5**). The two clusters with the highest proportion (>75%) of US ancestral birth origin (“Northwest Europe 1” and “Northwest Europe 2”) have ∼4.5% of UK ancestral origins. The Central European cluster and the Irish cluster both have 66.1% and 68.5% of US ancestral origins, respectively. In contrast, the US makes up only 62.2% and 34.5% of ancestral birth origin for the clusters of Southern Europeans and Eastern Europeans, respectively.

**Figure 5.**
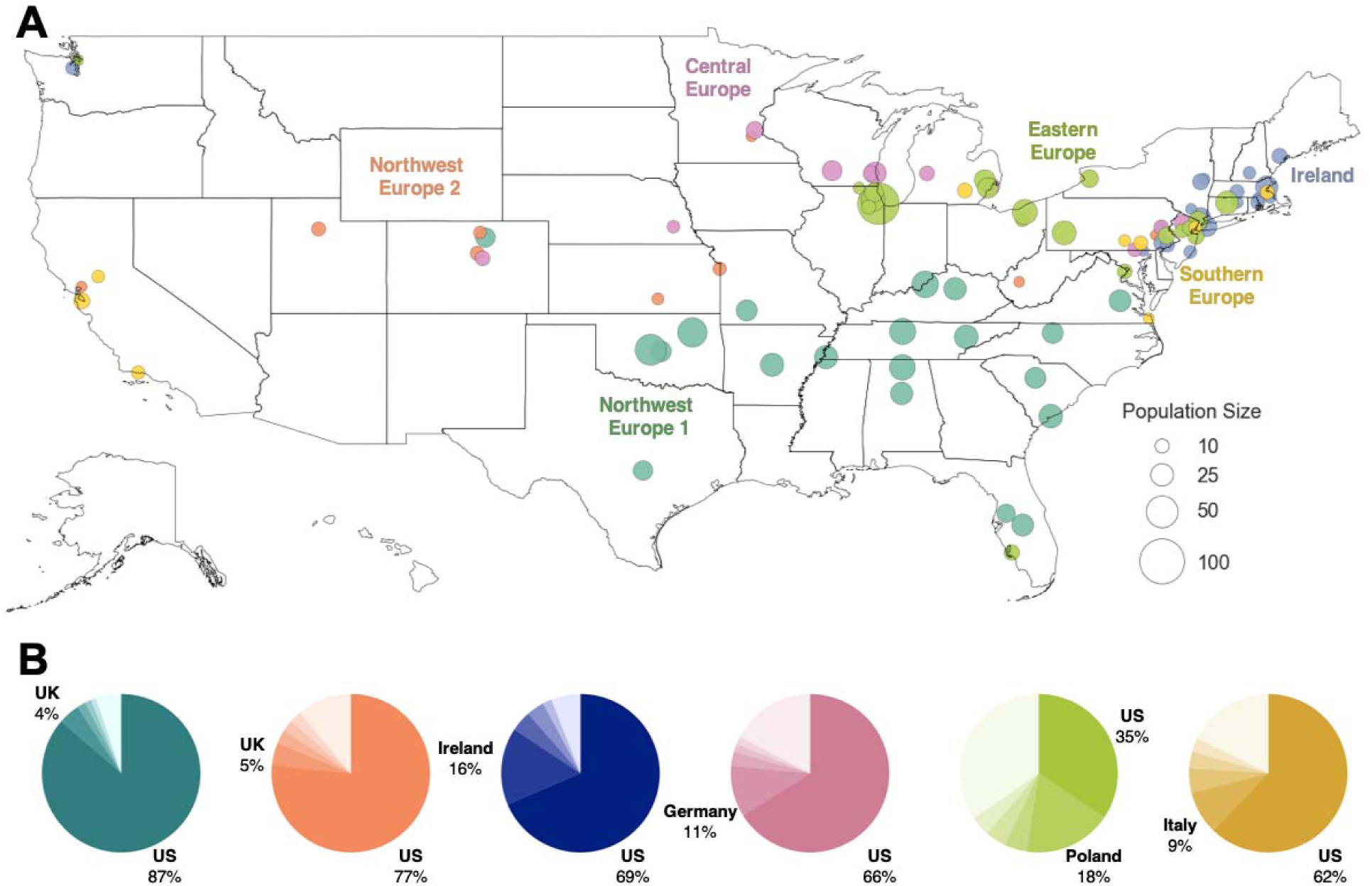
Distribution of European American Haplotype Clusters. (A) Geographic distributions of haplotype clusters that correspond to regional European ancestries. Each dot represents a county containing present-day individuals. The size of the dot represents the number of individuals of the particular cluster in that county. For each cluster, the top 20 locations with the highest odds ratio are shown. Maps showing the full distribution for each cluster can be found in the supplement (Figure S11). (B) Ancestral birth origin proportions for each cluster in (A). Only individuals with complete pedigree annotations, up to grandparent level, are included. For each chart, only the top five birth origins are shown as individual proportions; the remaining birth origins are aggregated into one slice (lightest color).

Unlike the larger European clusters, the smaller European clusters reflect the structure of recent immigrants and genetically isolated populations, recapitulating earlier findings.^8^ The geographic distributions of these subpopulations are more concentrated, and their ancestral birth origin proportions are overrepresented by specific countries and ethnicities (**Figure 6**). Specifically, Finns and Scandinavians are abundant in the Upper Midwest and Washington; French Canadians are found in the Northeast; Acadians are present in the Northeast and Louisiana; and Italians, Greeks, Ashkenazi Jews, and Admixed Jews are mostly located in the metropolitan area of New York City. Of the European clusters, median cumulative IBD sharing and cROH lengths are highest amongst Ashkenazi Jews (31.8cM and 11.3 Mb, respectively; **Table 1**), reflective of past founding events and endogamy.^17, 39^ The two Jewish-related clusters were identified using self-reported ancestral ethnicity data rather than birth origin data, since Jewish ancestry is not specific to any single location. Jewish ancestry, particularly Ashkenazi Jewish ancestry, was more consistently reported on both sides of the family in the larger Jewish cluster (“Ashkenazi Jewish”), suggesting that individuals are more admixed in the smaller cluster (“Admixed Jewish”).

**Figure 6.**
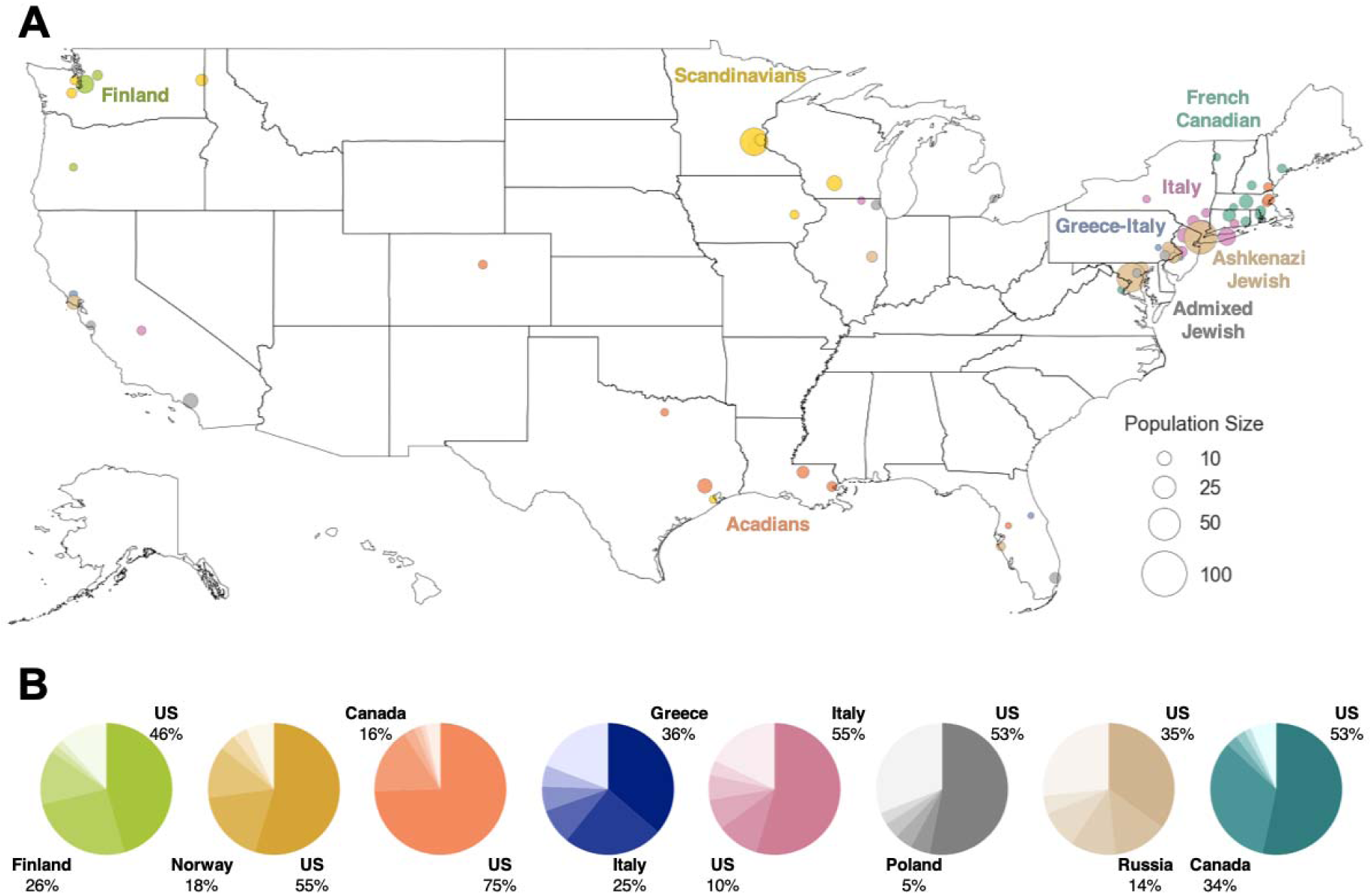
Distribution of Genetically Differentiated European American Haplotype Clusters. (A) Geographic distribution of present-day individuals in clusters corresponding to European populations that are more genetically isolated, displayed similarly to Figure 5A. For clarity, the top ten locations with the highest odds ratio are shown for each cluster. (B) Ancestral birth origin proportions for each cluster in (A). Only individuals with complete pedigree annotations, up to grandparent level, are shown. For each chart, only the top five birth origins are shown as individual proportions; the remaining birth origins are aggregated into one slice (lightest color).

We inferred two haplotype clusters of African Americans separated along a north-south cline, recapitulating the EEMS migration barrier inference. One cluster is primarily distributed amongst the northern and western states (“African Americans North”) while the other is distributed amongst the states southeast of the Appalachian Mountains (“African Americans South”) (**Figure S9**). The proportion of US birth origin is higher in the northern cluster than the southern cluster, further evidence of isolation by distance amongst African Americans in the north.^7^ These two clusters share similar cROH lengths but differ in admixture proportions and median IBD sharing, pointing to a cluster with consistent African American ancestors and a cluster with more admixed ancestors. Median IBD sharing is higher amongst African Americans in the south (median IBD = 19.6 cM, median cROH = 3.3 Mb) than in the north (median = 15.9 cM; **Table 1**) while the average proportion of African ancestry is higher in the northern cluster than the southern cluster.

Four of the clusters reflect recent immigrants from Asia (**Figure S10**), which grew rapidly in the mid-20th Century after the elimination of national origin quotas.^40^ The recency of immigration among these clusters is supported by the observation that fewer than 30% of grandparents were born in the US. Geographically, individuals in these clusters primarily reside in major cities. East Asians predominantly inhabit the metropolitan areas of the West and Northeast (OR > 2), Southeast Asians are enriched in the West (OR > 2.5), and South Asians are strongly associated with the Northeast (OR > 2.5). Despite its small size, the cluster of Greater Middle East individuals reflects many of the known demographic patterns of Arab Americans, as individuals in this cluster are primarily of Lebanese origin and are distributed in the Northeast as well as metropolitan Detroit. cROH lengths are particularly long for South Asians (median cROH = 10.3 cM), Southeast Asians (median cROH = 7.8 cM), and Middle Easterners (median cROH = 8.2 cM), potentially reflecting inbreeding patterns found in their ancestral regions.^41^

## Discussion

As the US population is becoming increasingly diverse, genomic studies are simultaneously growing in scale and relevance; to increase scientific and ethical parity, these studies must move beyond the current practice of evaluating genetically homogenous groups in isolation.^30, 42^ Here, we provide an integrative framework for analyzing population structure in ancestrally heterogeneous individuals. Our comprehensive approach has allowed us to capture spatial patterns of gene flow within and between subpopulations that are difficult to infer from a single method alone. For example, while EEMS enabled us to examine genetic similarity at a finer scale than previous studies and identify genetic differentiation within a state, EEMS can only compare neighboring demes and does not directly evaluate the genetic similarity of geographically distant individuals. Haplotype clustering, on the other hand, can identify population structures over long distances, but it does not measure genetic similarity with respect to geography. Since individuals are exclusively assigned to a single cluster, information regarding admixture, especially between neighboring clusters, are lost during haplotype clustering. An integrative approach can thus enable greater insights into populations with complex histories, as well as populations typically overlooked in previous studies such as Asian Americans.

The genetic structure and history of Hispanic/Latino populations is particularly complex due to many historical migration and admixture events.^4, 9^ This complexity is reflected in the variable migration rates across the country and the large variations in admixture proportions within and between subpopulations. While prior analysis of Hispanics/Latinos in the US found differences in ancestry proportions aggregated at the state level,^9^ we demonstrate that considerable differences in genetic ancestry also exist within a state. For example, two distinct clusters— Puerto Rico and Hispanics/Latinos—are found in Florida with the Puerto Rico cluster having higher average African ancestry proportions than the Hispanics/Latinos cluster (9.0% vs 2.5%, respectively). EEMS also enabled direct measures of genetic similarity within states and between subpopulations. While the mean ancestry proportions are similar between the New Mexican cluster and the Texan cluster, individuals in northern New Mexico are more genetically differentiated than individuals in southern New Mexico, as indicated by the migration barrier. The individuals in northern New Mexico are likely *Nuevomexicanos*, descendants of Spanish colonial settlers, while those in the south are more genetically similar to Hispanic/Latino individuals in central Texas, likely because they share a common ancestral origin (i.e. Mexico). We also built upon the use of pedigree annotation^43^ by quantifying ancestral origins to better understand the differences in genetic ancestry between subpopulations. For example, in the Hispanics/Latino in California cluster, the mean proportion of European ancestry is smaller when compared to the New Mexican and Texan clusters, reflecting the smaller proportion of US ancestral origin.

The demographic history of African Americans is characterized by large-scale migration and admixture, primarily due to the transatlantic slave trade and racial segregation.^34, 44^ The patterns of genetic ancestry and relatedness between states and regions of the US reflect these events.^3, 7, 9^ Our results show, at a finer scale, the barriers to migration and gene flow, particularly along the Appalachian Mountains. This migration barrier overlaps with the boundary between slave states and free states, as well as the boundary between states that enacted laws enforcing racial segregation and states that forbade segregation. The north-south separation of two African American clusters further emphasize this divide. The African Americans South cluster contains more recent ancestors from outside the US, particularly from the Caribbean, than the African Americans North cluster. These insights further emphasize the impact of historical migration and socioeconomic divide on the present-day patterns of genetic relatedness among African Americans.

Despite account for over 5% of the US population, individuals with Asian ancestries are underrepresented in US population genetics studies, hindering the ability of prior studies to investigate of their ancestry.^43^ Our analyses of these individuals therefore provide new insights into their genetic structure. Many of these individuals are descendants of recent immigrants, as indicated by the high proportions of non-US grandparental ancestral origin; therefore, they likely reflect the population of their ancestral region. The genetic structure of these individuals is particularly diverse. Using fineSTRUCTURE, genetic differentiation was found between East Asian and South Asian individuals of different ancestral origin as well as between individuals with the same ancestral origin. At the same time, longer cROH was observed in the Southeast Asia, South Asia, and Greater Middle East haplotype clusters, likely reflecting consanguinity or endogamy patterns in their ancestral countries. For example, the long cROH in South Asians may reflect endogamy related to the caste system in India, while similar patterns among the Middle Eastern and Southeast Asian clusters may be capturing consanguineous marriage practices in those regions.^31, 45, 46^ Understanding population genetic structure and patterns of homozygosity are important in determining the genetic profile of diseases within subpopulations, especially since these recent immigrants are becoming less similar to those in their ancestral countries due to outbreeding, admixture and population growth.^47, 48^ A populations mix, heterozygosity increases and allele frequencies change. This, in turn, can alter the prevalence of certain diseases, particularly rare recessive disorders which are often higher in populations with increased homozygosity.^49^ At the same time, changes in allele frequency can also reduce the accuracy of genetic predictors of complex traits (i.e. polygenic risk scores), especially if the prediction model was built using a homogeneous cohort of individuals from a divergent ancestry.^42^

Population history in the US is best characterized among individuals of European descent. Genetic diversity tends to be highest in more densely populated regions, likely due to multiple populations living in the same place. Many of the European subpopulations we identified are similar to those previously found—e.g., French Canadians, Acadians, Scandinavians, and Ashkenazi Jews (**Supplemental Discussion**).^8^ The geographic distribution of these subpopulations, particularly those that are more genetically diverged, overlap in the metropolitan areas of the Northeast, Midwest, and California.

The precision of population labels assigned to clusters of individuals is a function of demographic complexity and sample size. For example, Finnish ancestry is clearly European but genetically distinct from several other European populations due to historical bottlenecks, making this ancestry cluster relatively easily separable. By contrast, most Americans of European descent have heterogeneous ancestors from several northwestern European countries who have admixed over time, resulting in relatively evenly distributed ancestry overlapping that of present-day Europeans from multiple primarily northwestern countries. Additionally, while we identify and describe some substantial structure among Hispanic/Latino populations, considerably more is likely to exist and remains to be learned from larger and more diverse future studies. Similarly, sub-regional resolution into the ancestry of recent Asian immigrants to the US has been relatively limited in population genetics studies, and the structure of this immigration will be learned from larger future studies. The accuracy of self-reported birth records and variable granularity of geopolitical boundaries also provide additional considerations regarding the precision of population labels.

In addition to being of anthropological interest, understanding fine-scale human history and its role in shaping genetic variation is also important for interpreting the genetic basis of biomedical traits. The emergence of biobank-scale genomic data is enabling the imputation of pedigree structure regardless of whether some relatives have contributed DNA,^50^ greater insights into the impact of fine-scale population structure on genetic associations with disease,^10, 13, 22, 42, 51^ and population-based screening for individuals with serious genetic and health-related associations.^52^ As participation in genetic studies increases in the US, for example with the All of Us Research Program or with direct-to-consumer genetic tests that an estimated 26 million people have taken (Regalado, n.d.), so does the need for inferring more granular demographic histories in diverse study cohorts. Understanding such structure is important to account for stratification, prevent the overgeneralization of potentially confounded results, and avoid exacerbating existing Eurocentric study biases.^42, 53–55^ This study demonstrates how genetic data can be coupled with geographic and birth origin data to reconstruct such demographic histories, particularly in a large and heterogeneous population.

## Materials and Methods

### Human Subjects

The Genographic Project and Geno 2.0 Project received full approval from the Social and Behavioral Sciences Institutional Review Board (IRB) at the University of Pennsylvania Office of Regulatory Affairs on April 12, 2005. The IRB operates in compliance with applicable laws, regulations, and ethical standards necessary for research involving human participants. All DNA samples included in this study came from customers of the National Geographic Genographic Project, who have consented to have their results used in scientific research. Genographic Project participants would first order a DNA Ancestry Kit through the Genographic Project website, provide a saliva sample at home, mix the saliva sample with a stabilization buffer solution, and return via postal mail. DNA is then extracted from the saliva sample and genome-wide genotyping was performed (Genotyping and Quality Control).

In addition to providing a DNA sample, participants also provided geographic location (postal code), and, optionally, family history information in the form of ancestral birth origin and ethnicity (up to grandparental level) when they registered on the Genographic Project website to track and access the results of their DNA sample. All data was deidentified prior to research access. We limited our study to those individuals who provided valid geographic location in the United States. Approximately 75% of individuals provided complete pedigrees and family history data (Supplemental Materials and Methods).

### Genotyping and Quality Control

Participants of the Genographic project were sequenced with the GenoChip array,^25^ an Illumina iSelect HD custom genotyping bead array with approximately 150,000 Ancestry Informative Markers. Raw genotype data was quality controlled (QC) using PLINK v1.90b3.39.^56^ We filtered to keep samples with ≤ 0.1 missingness, sites with = 0.0 missingness, and MAF ≥ 0.05. A total of 32,589 individuals and 108,003 SNPs passed quality control.

### Ancestry Reference Panels

We leveraged a variety of reference populations to help better infer and interpret the genetic ancestry, admixture proportions, and population structure in the Genographic cohort. Data from the 1000 Genomes Project was used to help identify genetic ancestry and estimate admixture proportions.^1^ 108,003 SNPs were shared between the Genographic samples and the 1000 Genome Project samples. We also used data from the Population Reference Sample (POPRES) to help understand the population structure of individuals with European ancestry in the Genographic cohort.^57^ All analysis with the POPRES data was limited to the 46,710 SNPs that are shared between the two datasets. We also leveraged recently released sequence data for the Human Genome Diversity Project (HGDP) to expand the available set of ancestral populations from Asia.^58^ All analyses using the HGDP data was performed using the 105,944 SNPs shared between the samples in Genographic Project and HGDP.

### Principal Component Analysis

We performed principal component analysis (PCA) on the quality-controlled samples using FlashPCA version 2.0.^27^ For PCA analysis of all Genographic individuals, we leveraged the genotypes of all 2,504 individuals from the 1000 Genomes Project as reference samples. We first computed PCs across the 108,003 shared sites for 1000 Genome Project individuals. We then projected the Genographic individuals on the same principal component space using the flag: --project.

For PCA analyses of East Asian and South Asian populations, we used samples from 1000 Genome Project that correspond to the East Asian and South Asian super population. Similar to above, we first compute PCs for the 1000 Genome Project samples separately for East Asians and South Asians. We then projected East Asian and South Asian Genographic individuals onto the respective principal component space using: --project.

### Continental Ancestry Assignment

We assigned continental ancestry to each Genographic sample by using a random forest classifier. Using the PCs and known super population assignment (AFR=African, EUR=European, EAS=East Asian, AMR=American, and SAS=South Asian) from the 1000 Genome Project samples as training data, we applied the classifier to assign ancestry to each Genographic sample at 90% probability. We considered unassigned ancestries as “other” (OTH).

### Genetic Ancestry Proportion Estimation

We estimated admixture proportions using ADMIXTURE by first analyzing samples from the 1000 Genomes Project in unsupervised mode to learn allele frequencies.^26^ Then, we projected the learned allele frequencies onto the Genographic samples to obtain the admixture proportions using the flag: -P. We ran ADMIXTURE with K = 2-9 and chose K = 5 as the most stable representation.

For the analysis of East Asian and South Asian, we combined samples from HGDP and 1000 Genome Project together to build more expansive reference panels. Specifically, we combined 1000 Genome Project populations under the East Asian (EAS) super population label with HGDP samples that have the East Asian and Oceania region label; and 1000 Genome Project samples of South Asian super population label with Central South Asia label in HGDP. Similar to above, we first ran ADMIXTURE on the ancestral reference panels for East Asians and South Asians, separately. We then projected the learned allele frequencies onto the Genographic samples to obtain admixture proportions using the flag: -P. We tested a variety of clusters, K = 2-9, and chose K = 4 for East Asians and K = 3 for South Asians as the most stable representation.

### UMAP

We applied the Uniform Manifold Approximation and Projection (UMAP) method to visualize subcontinental structure.^28, 29^ We first combined the PCs of the Genographic samples and the 1000 Genome Project samples into one dataset. We then applied UMAP on the first 20 PCs from the joint dataset to produce a two-dimensional plot. We tested various parameter choices for UMAP and found that the default nearest neighbor value of 15 and the minimum distance values of 0.5 delivered the clearest result. Coloring of UMAP plots are described in the Supplemental Materials and Methods.

We further examined the subcontinental structure of Genographic individuals who were classified as Europeans with data from the Population Reference Sample (POPRES).^57^ Similar to the analyses with the 1000 Genome Project data, we performed dimensionality reduction with PCA and UMAP, keeping the same parameter values. Coloring of POPRES data was grouped by continental regions: Southeast Europeans = Croatia, Yugoslavia, Bosnia-Herzegovina, Serbia, Romania, Hungary, Albania, Macedonia; Central Europe = Switzerland, France, Germany, Germany, Swiss-Italian, Belgium, Swiss-French, Netherlands, Swiss-German; British Isle = Scotland, Ireland, United Kingdom; South Europe = Italy, Cyprus, Turkey, Greece; Iberian = Portugal, Spain; Eastern Europe = Austria, Czech Republic, Poland, Russia; Scandinavia = Sweden, Norway.

### Phasing and Haplotype Estimation

Genographic genotypes were phased with the Sanger Imputation Service using EAGLE2^59^ and the Haplotype Reference Consortium reference panel.^60^ No genotype imputation was performed.

### fineSTRUCTURE analysis

For classified East Asian individuals and South Asian individuals, we inferred clusters of unrelated individuals with shared ancestries by applying the fineSTRUCTURE framework v.4.0.1, a model-based approach to estimate patterns of haplotype similarity and identify clusters of discrete populations.^24^ We performed fineSTRUCTURE analysis separately for the two populations. The first part of the fineSTRUCTURE framework uses ChromoPainter to measure shared ancestry between individuals and estimate a coancestry matrix. This matrix is then used in fineSTRUCTURE’s clustering and tree-building algorithm to hierarchically clusters of individuals from fine levels of structuring to broader levels. We first applied ChromoPainter to phased genotypes to estimate the number of contiguous segments (chunks) shared and total amount of genome (in cM) shared between each pair of individuals within each population, as well as the normalization parameter (*c*). Using the coancestry matrix and normalized parameter, we then ran the fineSTRUCTURE with 2 million Markov Chain Monte Carlo (MCMC) iterations, of which 1 million are “burn-in” iterations, and every 2,000 iterations was sampled. Finally, we used fineSTRUCTURE to infer a hierarchical tree using 100,000 hill-climbing moves. We used the scripts accompanying the fineSTRUCTURE software as well as the *ape* package in R to visualize the coancestry matrix and dendrogram results.

To examine the properties of the inferred clusters, we sought to examine both structure both the broad-scale and fine-scale. There is no definitively correct level of the dendrogram to pick for examination. We examined clades at various levels of the tree and assessed broad structure at the levels in which clades had sufficient number of individuals (on average 50 or more samples). We further used a combination of PCA and analysis of ancestral origins to assess and define these clades. Some of the clusters are small but genetically distinct as evident by the branch length and height of the split (i.e. Girmitiyas, Bangladesh), and therefore, they were kept as separate clades.

Unlike traditional PCA, PCA using the coancestry matrix (i.e. chunk counts matrix) can better discern fine-scale population structure and provide greater interpretability.^24^ We performed PCA analysis on the chunk counts matrix using in the Python library *scikit-learn*. Individual markers are colored and labelled based on their respective grouping.

### Estimating Effective Migration Surfaces

We estimated migration and diversity relative to geographic distance using the estimating effective migration surfaces (EEMS) method for Genographic individuals that were classified under African, European, and Native American ancestries.^33^ We excluded East Asian and South Asian ancestries due to low sample size and density. We used unrelated individuals with available postal code data. We first computed pairwise genetic dissimilarities with the EEMS *bed2diffs* tool and then ran EEMS with *runeems_snps*, setting the number of demes to 250. Per the recommendation in the manual, we adjusted the variance for all proposed distributions of diversity, migration, and degree-of-freedom parameters such that all were accepted 10%-40% of the time. We increased the number of Markov chain Monte Carlo (MCMC) iterations until it converged.

### Haplotype Calling and Network Construction

We used IBDSeq version r1206 to generate shared identity-by-descent (IBD) segments from genotype data for all unrelated individuals.^61^ Unlike other IBD detection algorithms, IBDseq does not reply on phased genotype data and is less susceptible to switch errors in phasing that can cause erroneous haplotype breaks. We filter for IBD segments greater than 3cM. We removed segments that overlapped with long chromosomal regions (1 Mb) that had no SNPs across all unrelated individuals. These sites can result in false positives IBD sharing and likely correspond to centromeres and telomeres. We calculate the cumulative IBD sharing between individuals by summing the length of all shared IBD segments. We then constructed a haplotype network of unrelated individuals by defining vertices an individuals and edge weights between vertices as the cumulative IBD sharing between individuals. We filtered for edges with cumulative IBD sharing is ≥12 cM and ≤72 cM, as previously described.^8^

### Detection of IBD Clusters

While fineSTRUCTURE can identify population structure in admixed cohorts using haplotype similarity,^23^ fineSTRUCTURE does not scale to large sample sizes and is not recommended for samples >10,000.^24^ We therefore sought to identify clusters of related individuals in the haplotype network using the Louvain Method implemented in the igraph package for R. The Louvain Method is a greedy iterative algorithm that assigns vertices of a graph into clusters to optimize modularity (a measure of the density of edges within a community to edges between communities).^36^ The Louvain Method begins by first assigning each node as its own community and then adds node *i* to a neighbor community *j*. It then calculates the change in modularity and places *i* in the community with that maximizes modularity. The algorithm repeats this continuously and terminates when no vertices can be reassigned.

We partitioned the haplotype network into clusters by recursively applying the Louvain Method within subcommunities. At the highest level, we take the full, unpartitioned haplotype graph and identify a set of subcommunities. We isolate the vertices within each subcommunity, keeping only the edges between those vertices to create separate new networks. We then apply the Louvain Method to the new subgraphs. We repeat this process up to four levels. We combined subcommunities with low genetic divergence based on F_ST_ values of < 0.0001.

### Annotation of IBD Clusters

We used a combination of ancestral birth origins and self-reported ethnicities to discern demographic characteristics of each cluster. For each cluster, we quantified the proportion of each birth origin (i.e. country of origin) amongst all four grandparents, treating each grandparent’s origin equality. We use these proportions to inform population labels. Clusters in which a single non-US birth origin was in high proportions was labeled with that country. In cases where multiple non-US birth locations exists in approximately equally high proportions, we assigned a label representing the broader region (e.g. Eastern Europeans for Poland, Lithuania, Ukraine, and Slovakia; East Asia for Japan, China). For certain clusters, annotations could not be easily discerned by birth origin data. In these cases, we relied on self-reported ethnicities to label the clusters as these populations were found to be less associated with a non-US country (e.g. Ashkenazi Jews) or the population has resided in the US for generations (African Americans, Acadians).

### Runs of Homozygosity

We used PLINK v1.90b3.39 to infer runs of homozygosity with a window of 25 SNPs.^56^ We calculated the cumulative runs of homozygosity (cROH) size by summing the lengths of homozygous segments.

### Local Ancestry Inference

We inferred local ancestry with RFMix v1.5.4 for Genographic samples in clusters that were annotated as Hispanics/Latinos and African Americans.^37^ We used samples of African (LWK, MSL, GWD, YRI, ESN, ACB, and ASW; N = 661), European (CEU, GBR, FIN, IBS, and TSI; N = 503), and Native American (MXL, PUR, CLM, and PEL; N = 347) ancestry from the 1000 Genomes Project to build the reference panel for classifying genomic segments. We ran RFMix with the default minimum window size (0.2 centimorgans, cM) and a node size of 5 with the flags: -w 0.2, -n 5. We then collapsed the output of RFMix, which denotes the classified ancestry of each SNP for each individual, into local ancestry segments/tracts (in cM) for each individual. We then derived global ancestry proportions for each individual using that individual’s local ancestry tracts; we summed the length of local ancestry tracts for each ancestry (EUR, AFR, AMR) dividing by the total length of the genome to get the global proportion of each ancestry. Global ancestry proportions were visualized using the *python-ternary* package in Python (see Web Resources).

### Genetic Divergence

We computed weighted Weir-Cockerham F_ST_ estimates for each pair of haplotype clusters using PLINK v1.90b3.39.^56^ Using the distance matrix of F_ST_ values between clusters, we constructed an unrooted phylogenetic tree using the neighbor joining method implemented in *scikit-bio* (see Web Resources). We visualized the tree using Interactive Tree Of Life (see Web Resources).

## Data and Code Availability

Genotype data and associated metadata are available to researchers through an application process and data usage agreement. We encourage qualified researchers to email the Genographic team at National Geographic Society for information on and access to the Genographic database.

Custom scripts generated to analyze the data in this paper are available through GitHub (https://github.com/chengdai/genographic_ancestry).

## Acknowledgement

We thank the National Geographic Genographic Project participants who consented to research for making this study possible. We also thank Gregory Vilshansky for helping organize and manage the data for the Genographic Project.

This work was supported by funding from the National Institutes of Health (K99MH117229 to A.R.M.). C.L.D., M.M., R.T., and C.R. would also like to thank all the members of the MIT Senseable City Lab Consortium for supporting this research. M.G.V. acknowledges support from the National Geographic Society.

## Author Contributions

C.L.D. and A.R.M. designed the study, performed research, and wrote the manuscript. R.S.W. founded and formerly directed the Genographic Project. M.G.V. coordinated and supervised the Genographic Project. M.M.V., C.H.Y., and R.T. contributed to the data aggregation and data analysis. A.R.M., C.R. and M.J.D. supervised research. All authors reviewed the manuscript.

## Web Resources

Analysis scripts (https://github.com/chengdai/genographic_ancestry);

1000 Genome Project (https://www.internationalgenome.org);

Human Genome Diversity Project (https://www.hagsc.org/hgdp/);

POPRES: Population Reference Sample (https://www.ncbi.nlm.nih.gov/projects/gap/cgi-bin/study.cgi?study_id=phs000145.v4.p2);

Ancestry pipeline (https://github.com/armartin/ancestry_pipeline);

python-ternary (https://github.com/marcharper/python-ternary);

scikit-bio (http://scikit-bio.org/);

Interactive Tree of Life (https://itol.embl.de/);

