## Supplemental Text and Figures 1-11 for "Population histories of the United States revealed through fine-scale migration and haplotype analysis"

**Supplemental Materials and Methods**

**Self-reported Ancestral Birth Origin and Ethnicity**

As part of the registration process to track and access the results of their DNA sample on Genographic Project website, Genographic participants were given the option to report birth origin data and ethnicity data on themselves, their parents, and their grandparents. A total of 24,566 individuals (75.4%) provided complete data (i.e. no missing data for any ancestors), resulting in 171,962 pedigree records. All analysis using ancestral birth origin and ethnicity data was performed using data at the grandparent level. Birth origin data was recorded at the country level, with the exception of certain territories and regions being listed separately. Participants provided ancestral birth origin data by selecting from a list of countries for each ancestor. Ethnicity data was provided in the form of free text and was therefore not standardized across participants, making aggregating and comparing self-reported ethnicity data challenging.

It is important to note that ancestry, ethnicity, and race are all complex concepts that result from many factors, including appearance, culture, socioeconomics, geography, etc. Therefore, ancestry, ethnicity, and race are not directly comparable, and there are limitations to comparing genetic ancestry with data on race and ethnicity from the US Census. For example, population genetic studies often analyze Hispanic/Latino, European American, and African American individuals separately.^1,2^ The US Census, however, classifies race and Hispanics/Latinos origin to be two separate and distinct concepts; Hispanics/Latinos may be of any race.^3^ As such, comparing the proportion of genetically-labeled Hispanic/Latino individuals in the US with the proportion of people declaring Hispanic/Latino origin in US Census is invalid as the percent of Hispanic/Latino origin in the US Census should not be added to percentages for racial categories.^3^

**Family Relationship Inference**

We used KING v2.0 to identify the set of unrelated individuals within the Genographic dataset separated by at least two degrees of relatedness.^4^ 806 individuals had kinship coefficients greater than 0.0884 and were removed for downstream analysis using EEMS and haplotypes.

**Coloring of UMAP plots**

We colored the 1000 Genome Project samples in the UMAP plot based on their country level assignments (**Figure 1C left**) and visualized the Genographic samples by coloring each sample based on their ancestry proportions from ADMIXTURE (**Figure 1C right**). Specifically, the color (RGB value) of each Genographic sample is a linear combination of the sample’s admixture proportions and the RGB values of each ancestry’s color (EUR = red, AFR = yellow, NAM = green, EAS = blue, SAS = purple).

**Comparison of filtered and unfiltered haplotype network**

We evaluated two networks: one with filtering for minimum or maximum IBD sharing and one with pairs of individuals in which cumulative IBD sharing is ≥12 cM and ≤72 cM, similar to prior analysis.^2^ Clustering of haplotype networks resulted in a total of 25 clusters for the filtered network (≥12 cM and ≤72 cM). For the unfiltered network, we arrived at 32 clusters, 4 of which had less than 10 individuals and were removed from subsequent analyses. Annotations for the 25 clusters from the filtered network were found to be more interpretable than annotations for the 28 clusters from the unfiltered networks. Specifically, many of the clusters from the unfiltered networks exhibited similar proportions of ancestral origins or ethnicities and were difficult to differentiate (**Table S4 and S5**). Certain populations (e.g. Finns, Middle Easterners) found from the filtered network were also not identified from the unfiltered network. We therefore used the 25 clusters from the filtered network in downstream analyses.

**Supplemental Discussions**

The five Hispanic/Latino-related clusters we identified recapitulate the state-by-state differences of the Hispanics population as reported in the US Census.^3^ The presence of the Hispanics cluster and the Puerto Rican cluster in Florida are consistent with the large proportions of Hispanics/Latinos in Florida reporting Puerto Rican (20%) and Cuban (29%) origin in US Census. Similarly, the distribution of the Puerto Rican cluster around New York City is in line with the high proportions (31%) of Hispanics/Latinos reporting Puerto Rican origin in New York state. In Southwestern states, smaller proportions of Hispanics/Latinos reporting Central and South America origins are found in Arizona than neighboring California (3% in Arizona versus 10% in California) in the US Census, consistent with our ancestral birth origin data.

Variations in the proportion of African ancestry amongst African Americans in the Genographic Project are consistent with previous studies.^1,5^ However, the mean proportion of African ancestry is slightly lower, potentially due to sampling bias. The number of African Americans in the Genographic cohort is smaller than the that for European Americans and Hispanics/Latinos. This may impact the EEMS result as the barriers of migration that are present from Iowa to Utah and from Texas to New Mexico may be influenced by increased noise from the fewer African American individuals that are sampled or living there.

The European populations of the US are relatively homogenous, as demonstrated by the low genetic differentiation (F_ST_) between haplotype clusters and the relatively high migration rates across much of the country. The distribution of migration barriers largely corresponds to metropolitan regions, many of which are associated with numerous European subpopulations. The largest European haplotype clusters reflect broader regional ancestries, as corresponding birth origins are not clearly overrepresented in any particular European origin. The exception is the cluster of Irish individuals (“Ireland”). During the 19^th^ and early 20^th^ centuries, millions of Irish immigrants entered into the US, which experienced religious tensions and discrimination and resulting in high rates of in-group marriage amongst Irish individuals.^6^ Nonetheless, present-day Irish Americans remain genetically similar to other Europeans from the central and northwestern parts of Europe, as reflected in the low F_ST_ values between those subpopulations. F_ST_ values were highest amongst more genetically isolated Europeans, as well as more southern Europeans. Consistent with previous analysis,^2^ we identify clusters of Scandinavians, Finns, French Canadians, Acadians, Ashkenazi Jews, Italians, and Greeks. We also identify a second cluster with Jewish ancestry (“Admixed Jews”), but unlike the Ashkenazi Jewish cluster, median cROH and IBD lengths are substantially lower. In this cluster, self-reported ethnicity suggests admixture between Jewish and non-Jewish ancestry individuals, as Jewish-ancestry is typically present only on one side of the family.

**Supplementary Figures**

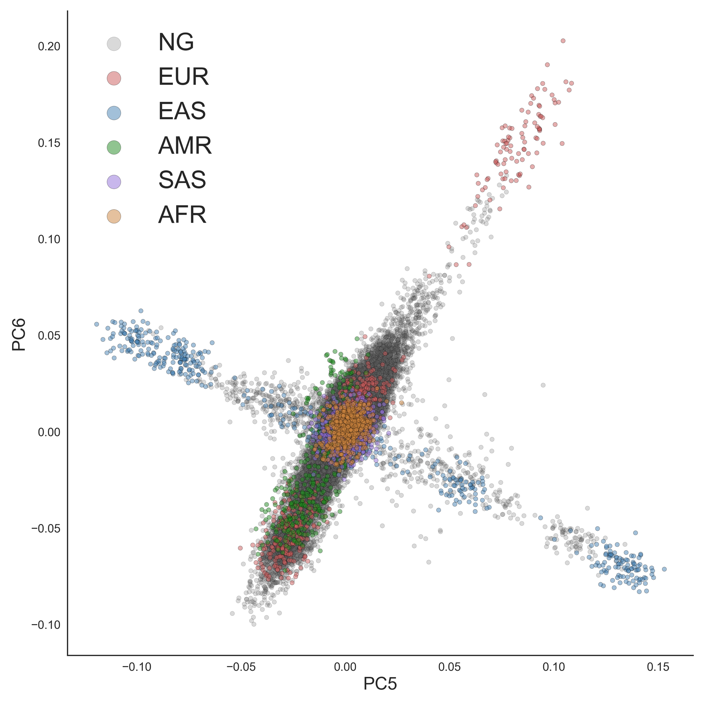

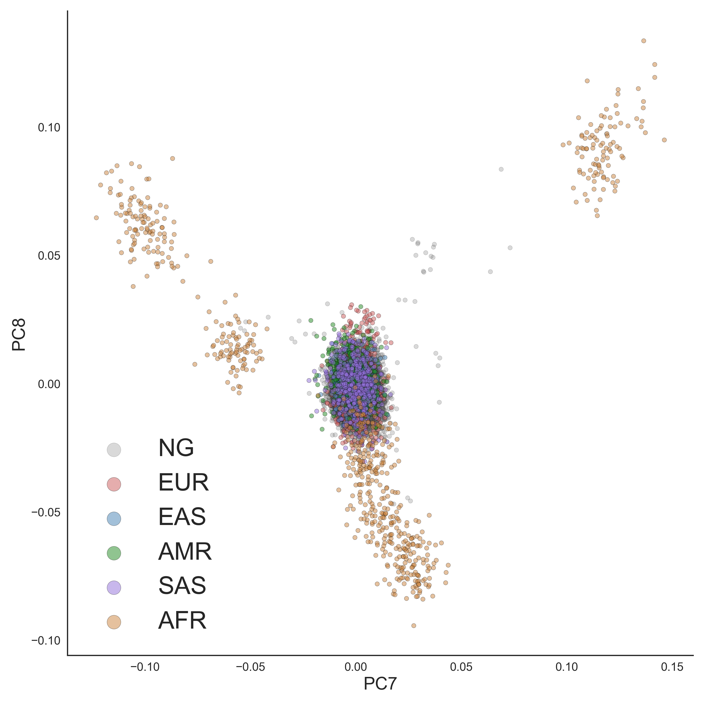

**Figure S1. Principal Component Analysis of 1000 Genome Project and Genographic samples**

PCA projects at for PC 5 and PC 6 (left); and for PC 7 and PC 8 (right).

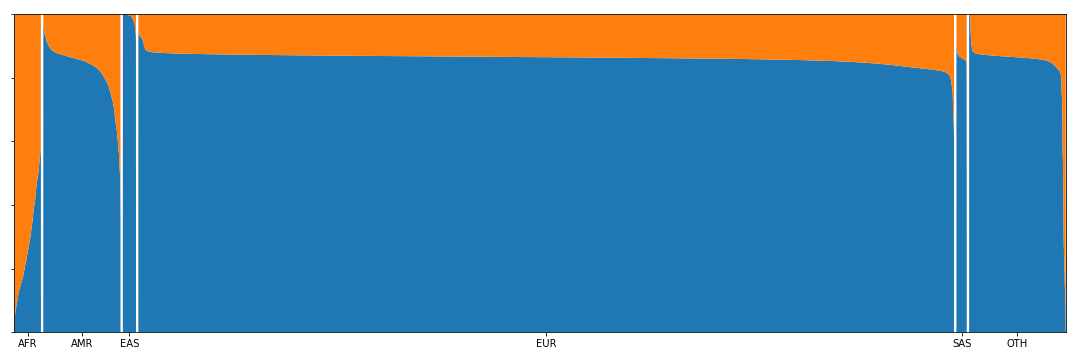

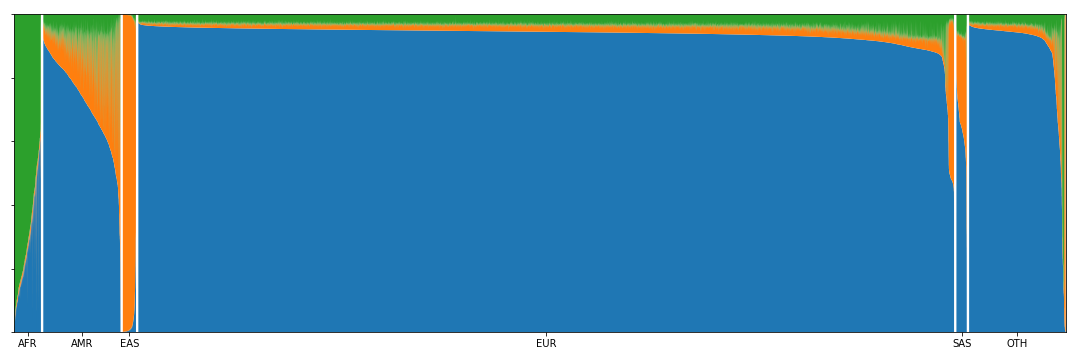

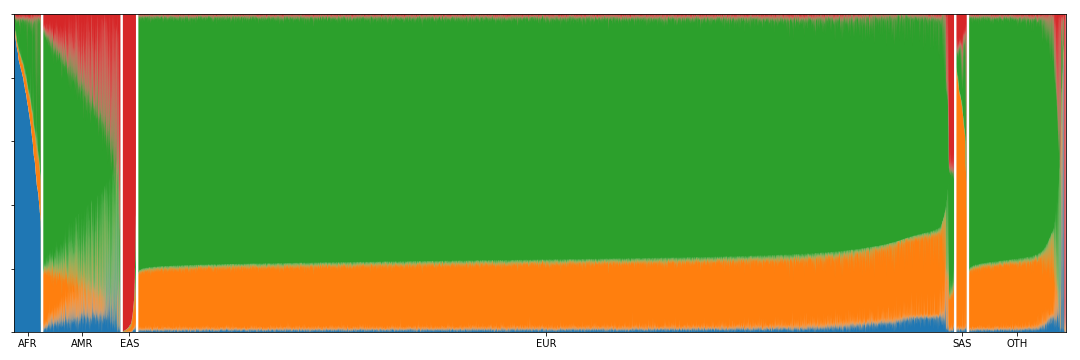

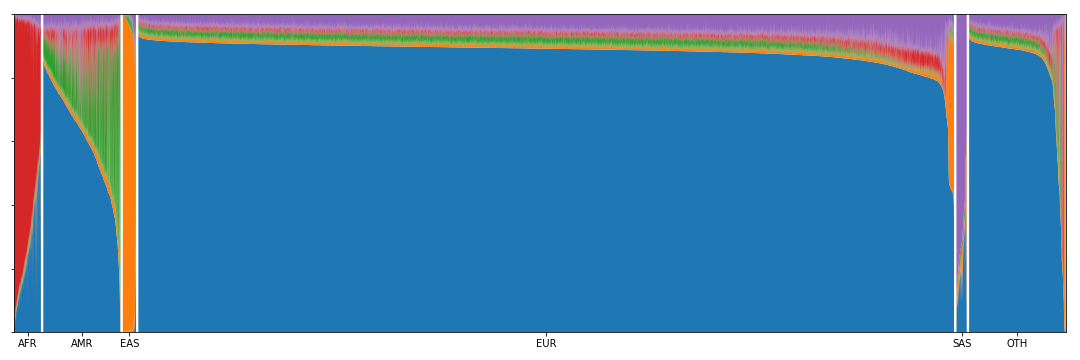

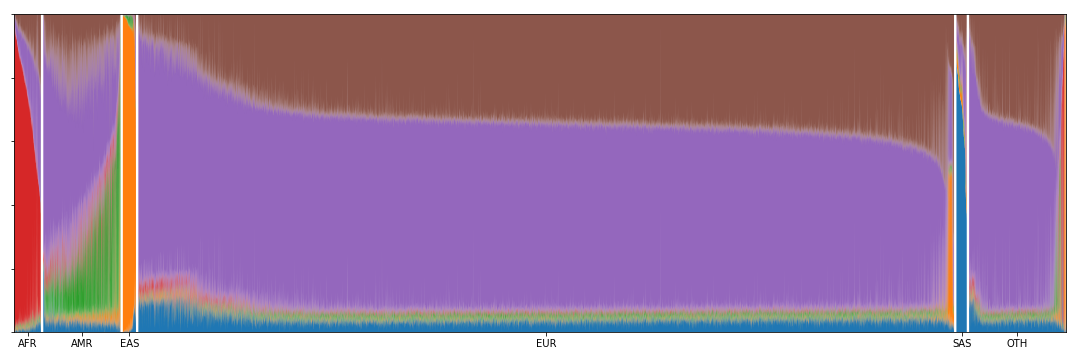

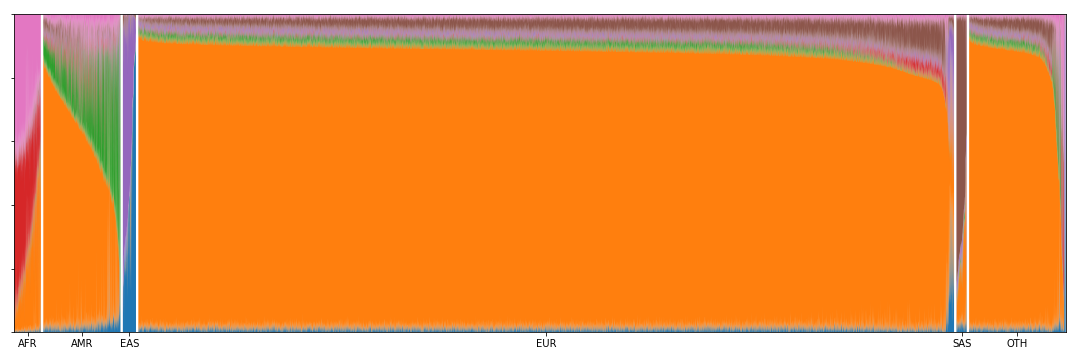

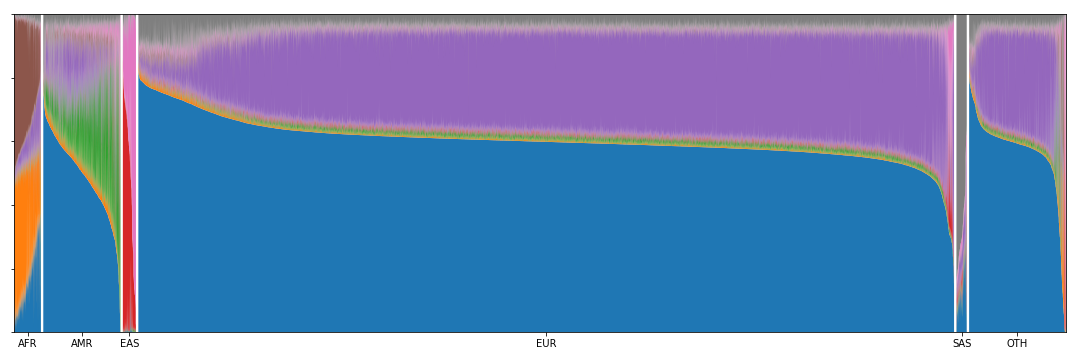

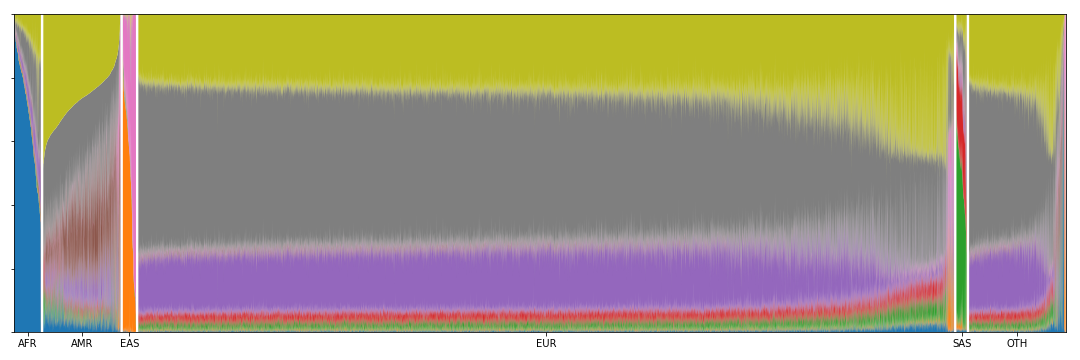

**Figure S2. ADMIXTURE from K = 2 to 9.**

ADMIXTURE analysis results for each K between 2 and 9 of U.S. individuals. Individuals were classified into continent level ancestry groups with a Random Forest model trained on the PCs from the 1000 Genome Project dataset.

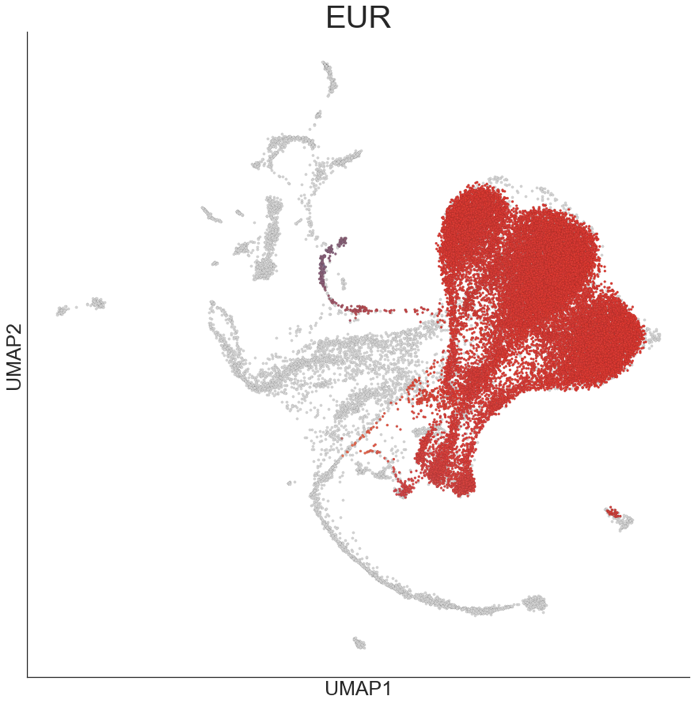

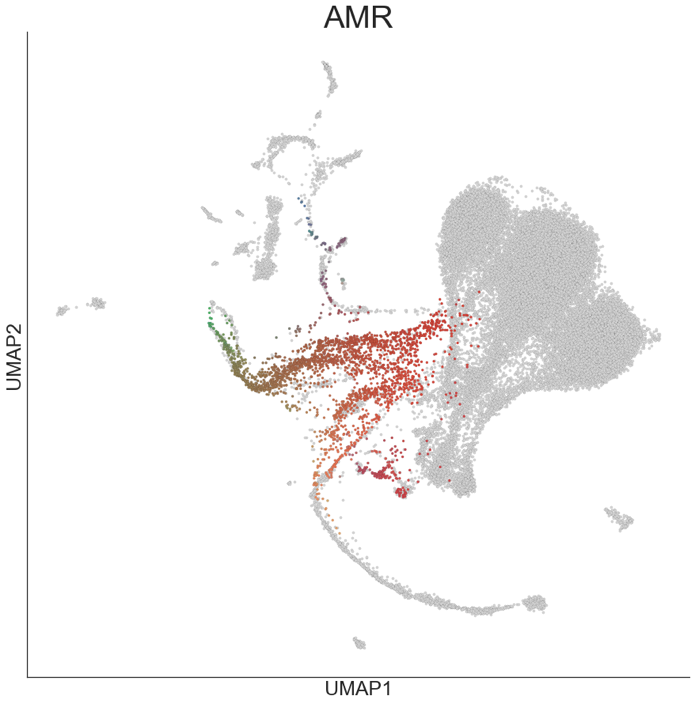

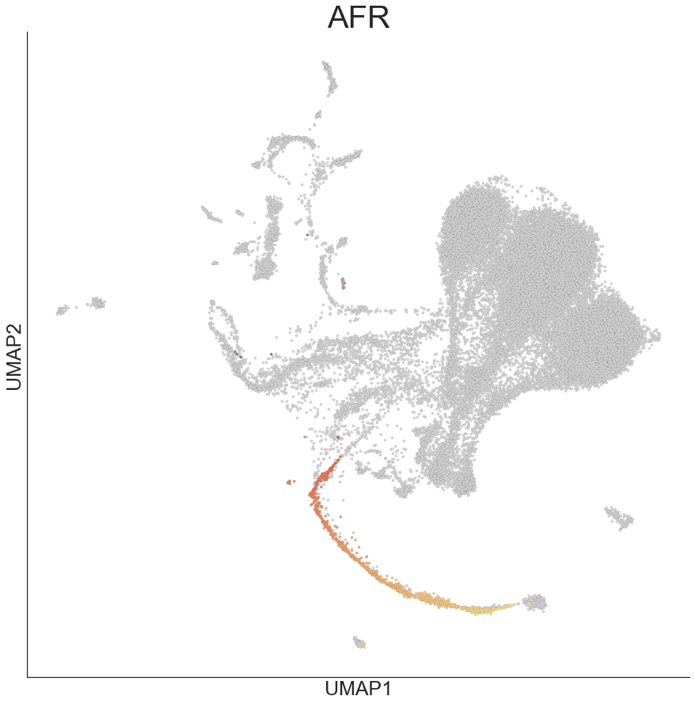

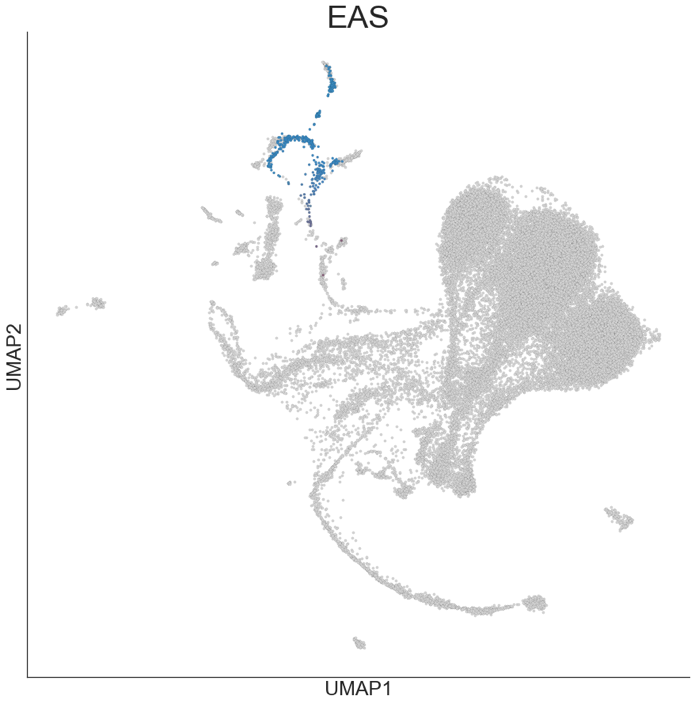

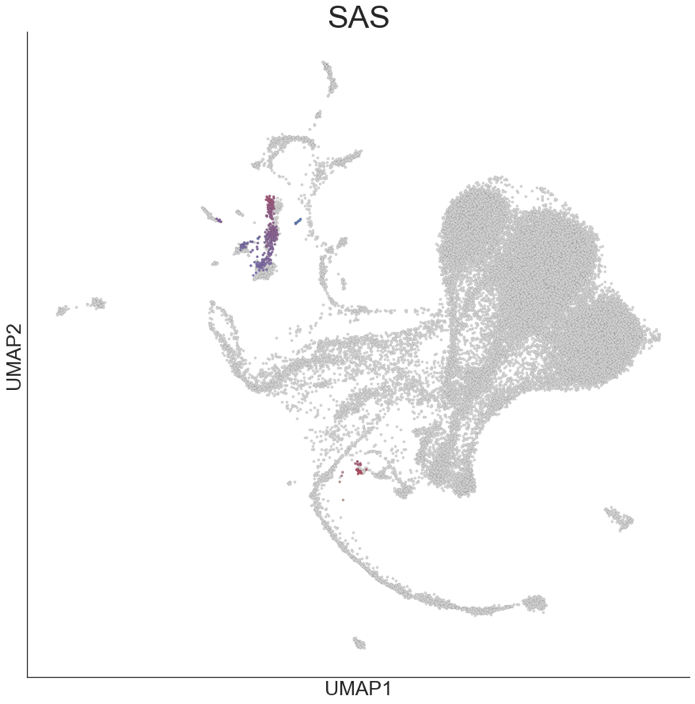

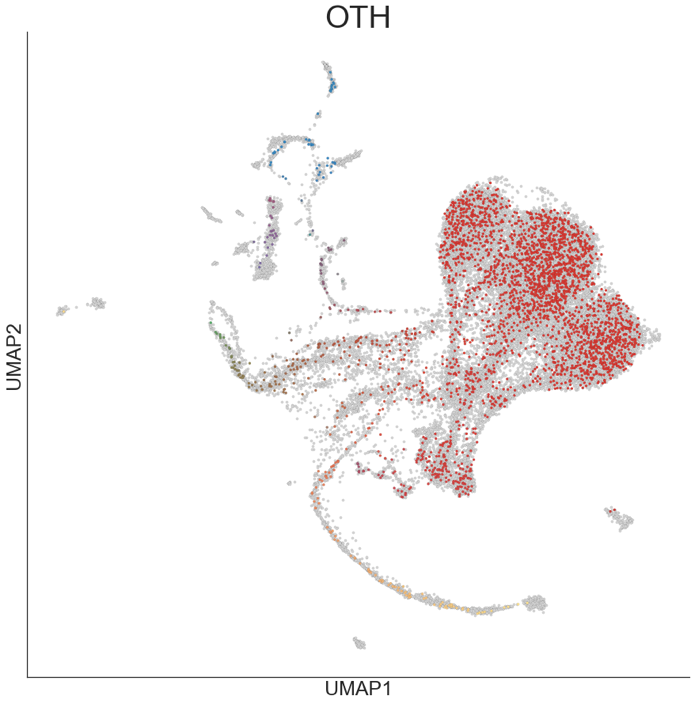

**Figure S3. Uniform Manifold Approximation and Projection (UMAP) of classified Genographic individuals**

UMAP projection of the first 20 PCs. Each dot represents one individual. Each plot represents the set of individuals classified at continental-level ancestry with the Random Forest model trained on the 1000 Genomes Project data. 1000 Genome Project individuals are colored in grey while U.S. individuals are colored based on their admixture proportions from ADMIXTURE. The color for each dot was calculated as a linear combination of each individuals admixture proportion and the RGB values for the colors assigned to each continental ancestry (EUR = red, AFR = yellow, NAM = green, EAS = blue, SAS = purple). Continental level ancestries are: EUR = European, AFR = African, NAM = Native American, EAS = East Asian, SAS = South Asian.

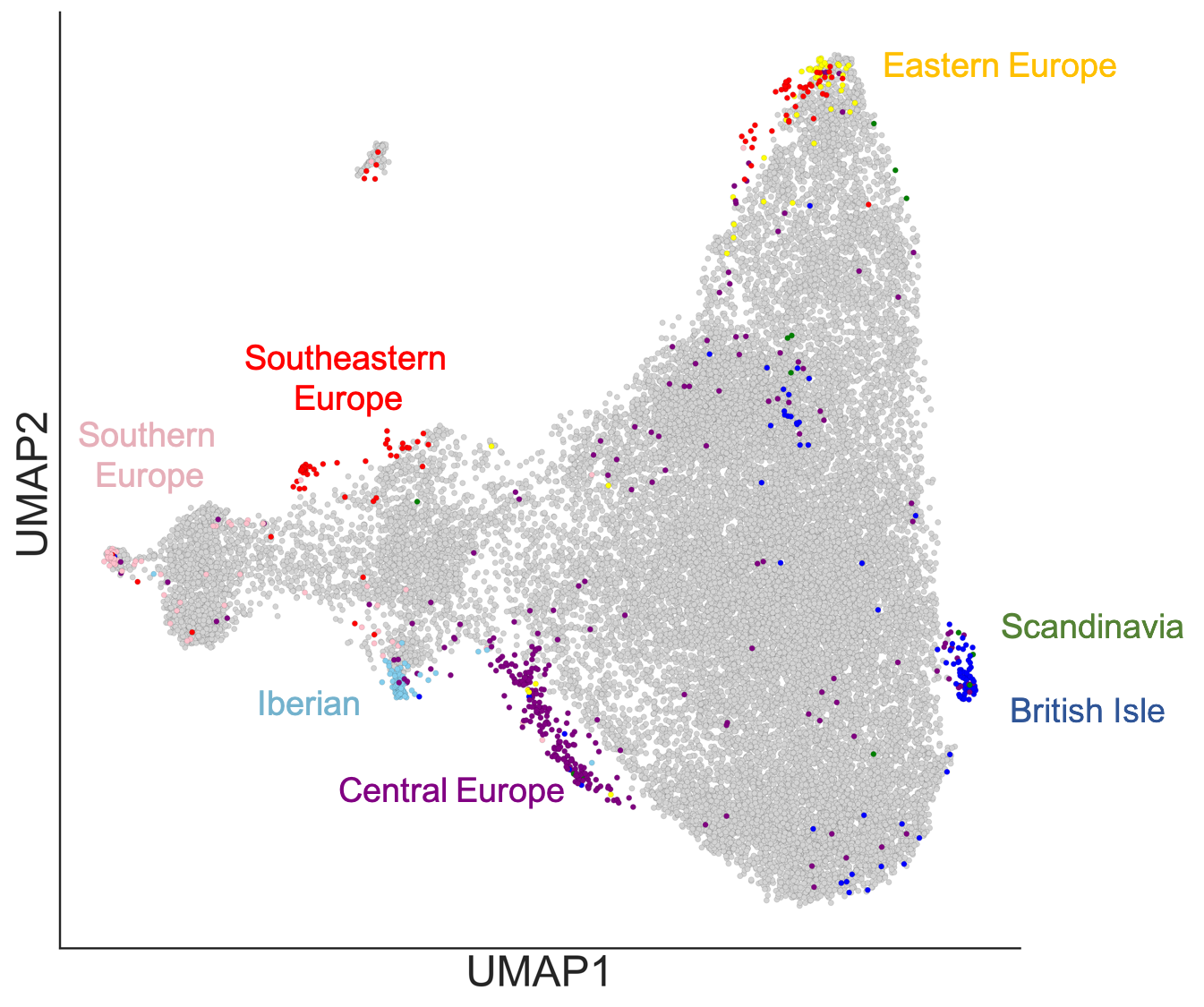

**Figure S4. UMAP of Classified Genographic European Americans and POPRES reference samples.**

UMAP projection of the first 20 PCs. PCs were calculated by first finding the PCs of the POPRES reference samples and then projecting the Random Forest classified Europeans in the Genographic cohort. Each dot represents one individual. Southeast Europeans = Croatia, Yugoslavia, Bosnia-Herzegovina, Serbia, Romania, Hungary, Albania, Macedonia; Central Europe = Switzerland, France, Germany, Germany, Swiss-Italian, Belgium, Swiss-French, Netherlands, Swiss-German; British Isle = Scotland, Ireland, United Kingdom; South Europe = Italy, Cyprus, Turkey, Greece; Iberian = Portugal, Spain; Eastern Europe = Austria, Czech Republic, Poland, Russia; Scandinavia = Sweden, Norway.

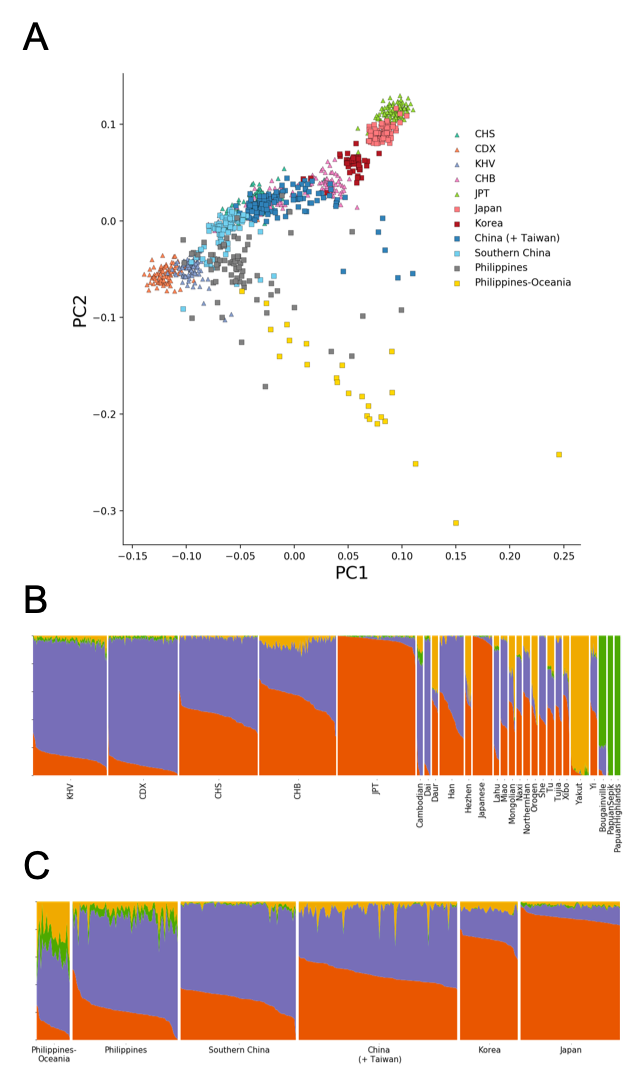

**Figure S5. PCA and ADMIXTURE analysis of East Asians**

**(A)** PCA analysis of classified unrelated East Asian Genographic individuals (plotted in squares) with East Asian samples from 1000 Genome Project (plotted in triangles). Genographic individuals are colored based on fineSTRUCTURE grouping (clade-level) while 1000 Genome Project Samples are colored based on super population.

**(B)** ADMIXTURE analysis of East Asian 1000 Genome Project samples (left five sections) and East Asia and Oceania HGDP samples (right 21 sections)

**(C)** ADMIXTURE analysis of classified East Asian Genographic individuals, grouped by fineSTRUCTURE clades.

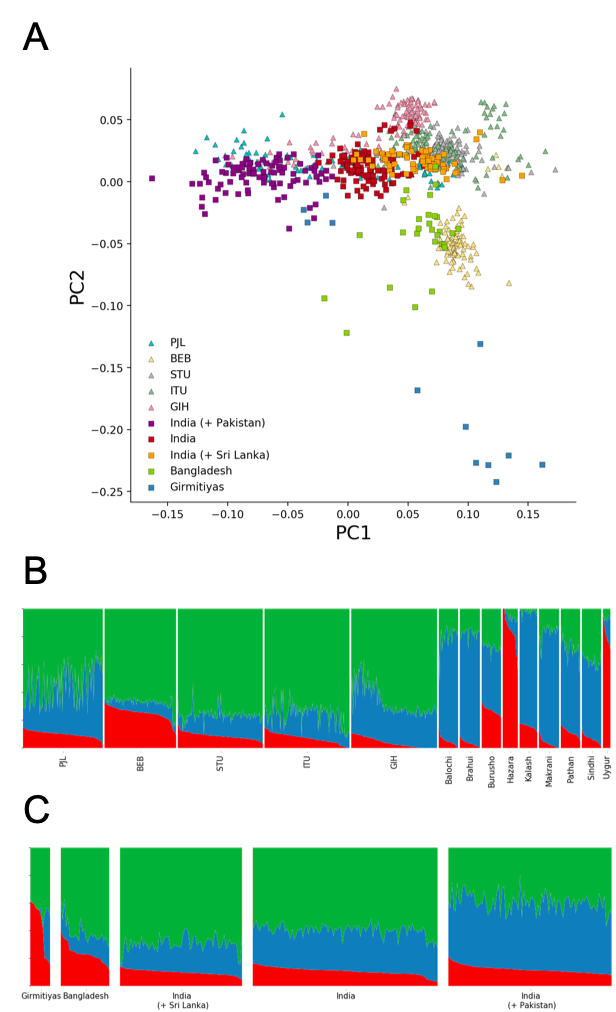

**Figure S6. PCA and ADMIXTURE analysis of South Asians**

**(A)** PCA analysis of classified unrelated South Asian Genographic individuals (plotted in squares) with South Asian samples from 1000 Genome Project (plotted in triangles). Genographic individuals are colored based on fineSTRUCTURE grouping (clade-level) while 1000 Genome Project Samples are colored based on super population.

**(B)** ADMIXTURE analysis of South Asian 1000 Genome Project samples (left five sections) and Central & South Asia HGDP samples (right nine sections)

**(C)** ADMIXTURE analysis of classified South Asian Genographic individuals, grouped by fineSTRUCTURE clades.

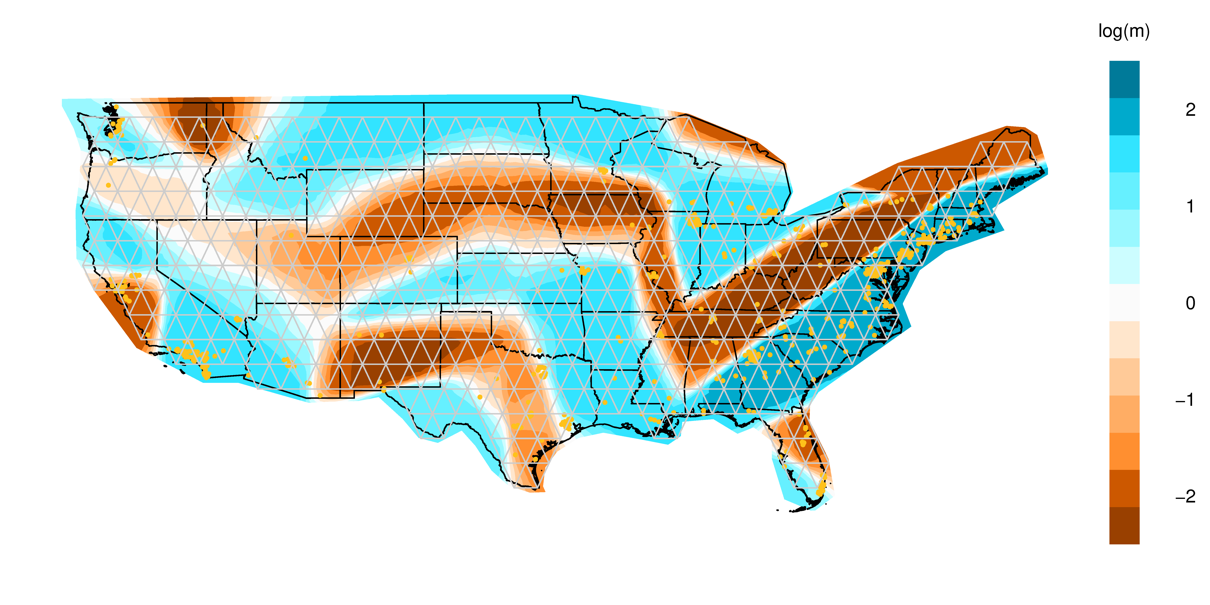

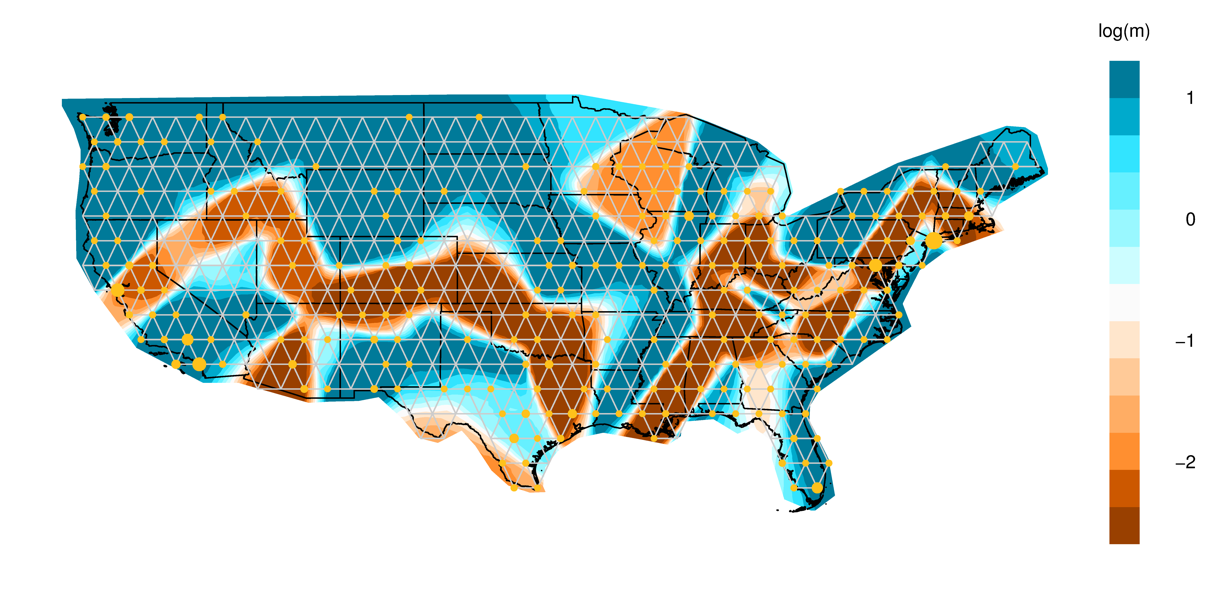

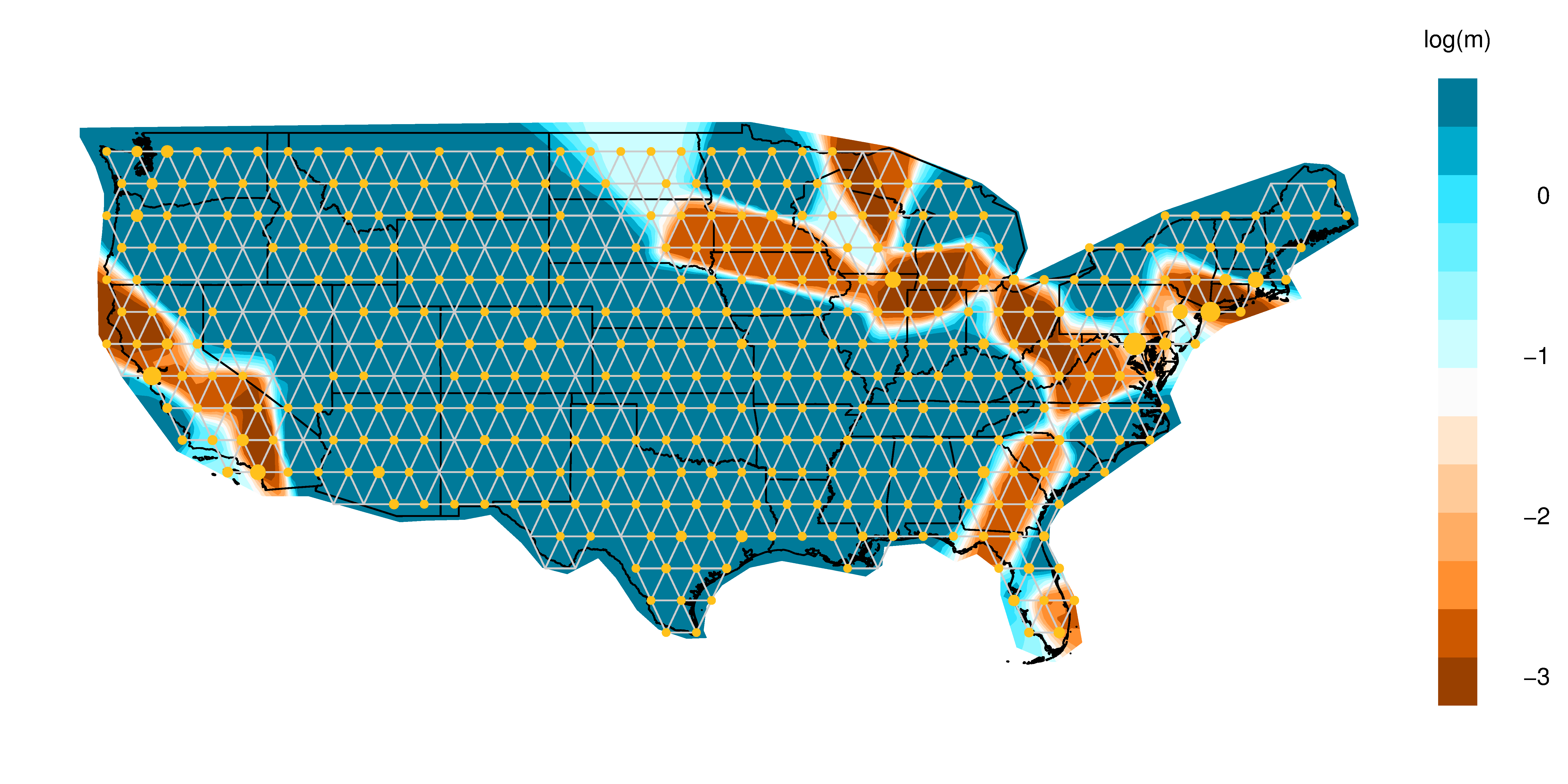

**Figure S7. Estimated Effective Migration Surfaces with demes.**

Migration rates inferred with EEMS for African Americans (top), Hispanics/Latinos (middle), and Europeans (bottom). Colors and values correspond to inferred rates, m, relative to the overall migration rate across the country. Shades of blue indicate logarithmically higher migration (i.e. log(m) = 1 represents effective migration that is tenfold faster than the average) while shades of orange indicate migration barriers. Each individual is snapped to a vertex, which is represented by yellow points. The size of points corresponds to the size of the subpopulation at the vertex.

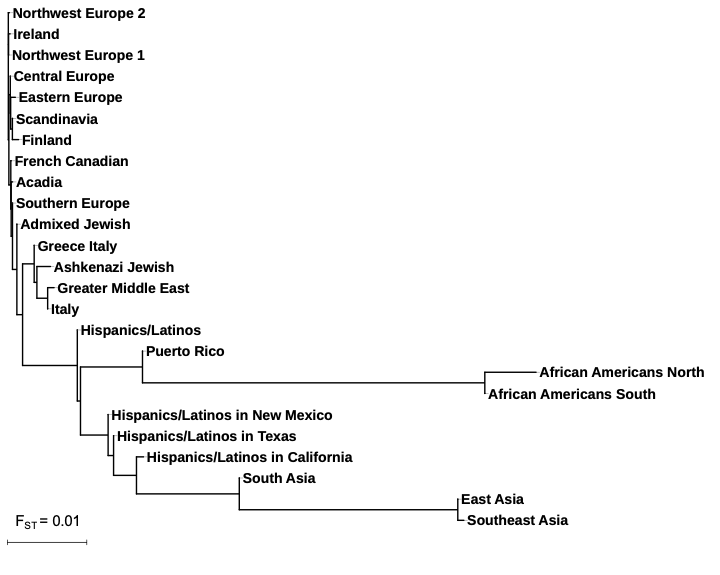

**Figure S8. Genetic differentiation of haplotype clusters**

Unrooted phylogenetic tree of haplotype clusters was constructed using the neighbor joining method with F_ST_ as genetic distance. Negative branch lengths were converted to zero.

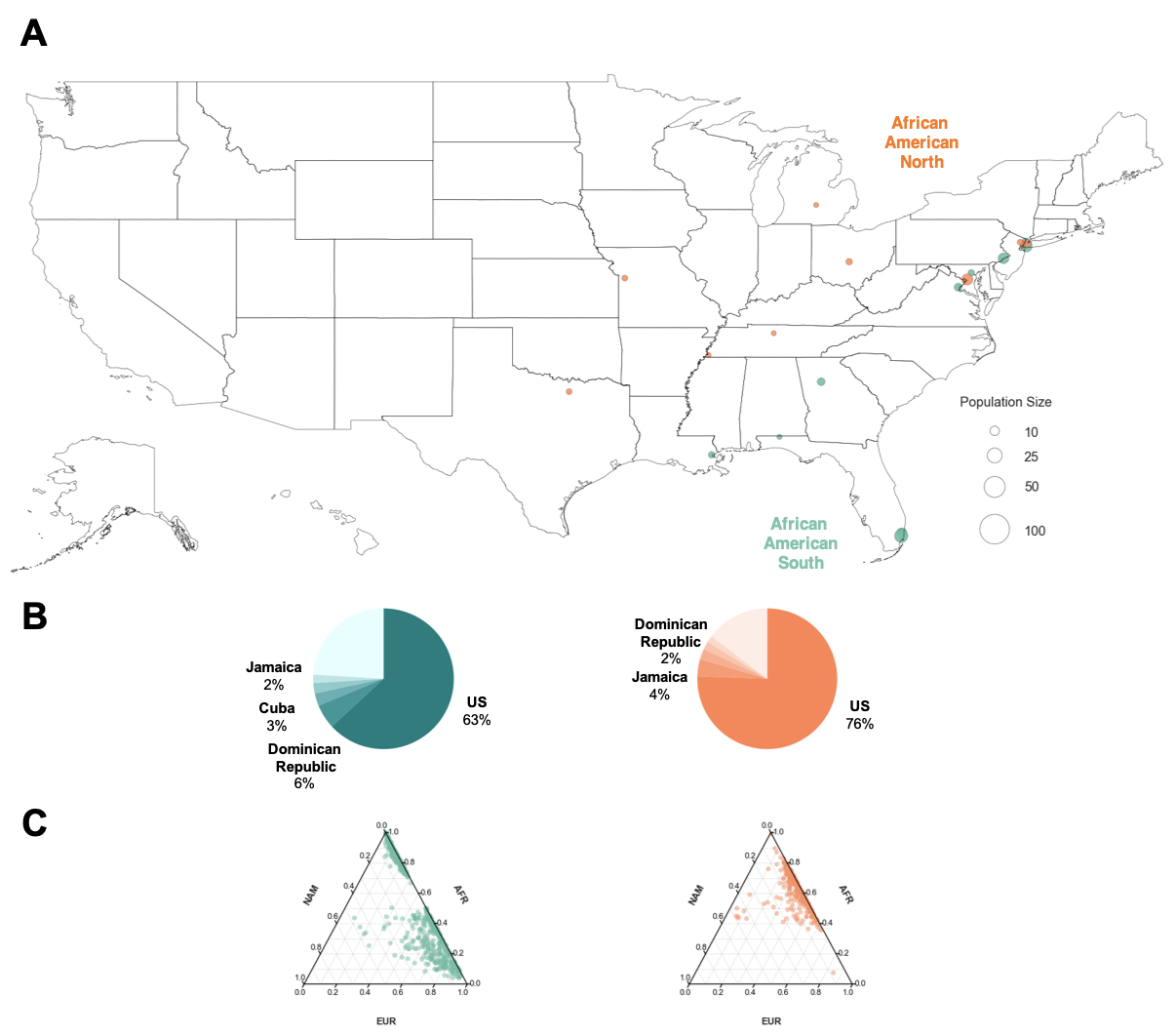

**Figure S9. Distribution of African American Haplotype Clusters**

**(A)** Map of haplotype clusters corresponding to Africans ancestries. Each county containing present-day individuals is represented by a dot. The top 10 locations with the highest odds ratio are shown for each cluster. Maps showing the full distribution for each cluster can be found in the supplement (**Figure S11**).

**(B)** Ancestral birth origin proportions for each cluster in (A). Only individuals with complete pedigree annotations, up to grandparent level, are shown.

**(C)** Ternary plots of ancestry proportions based on local ancestry inference for each haplotype cluster. Each dot represents one individual.

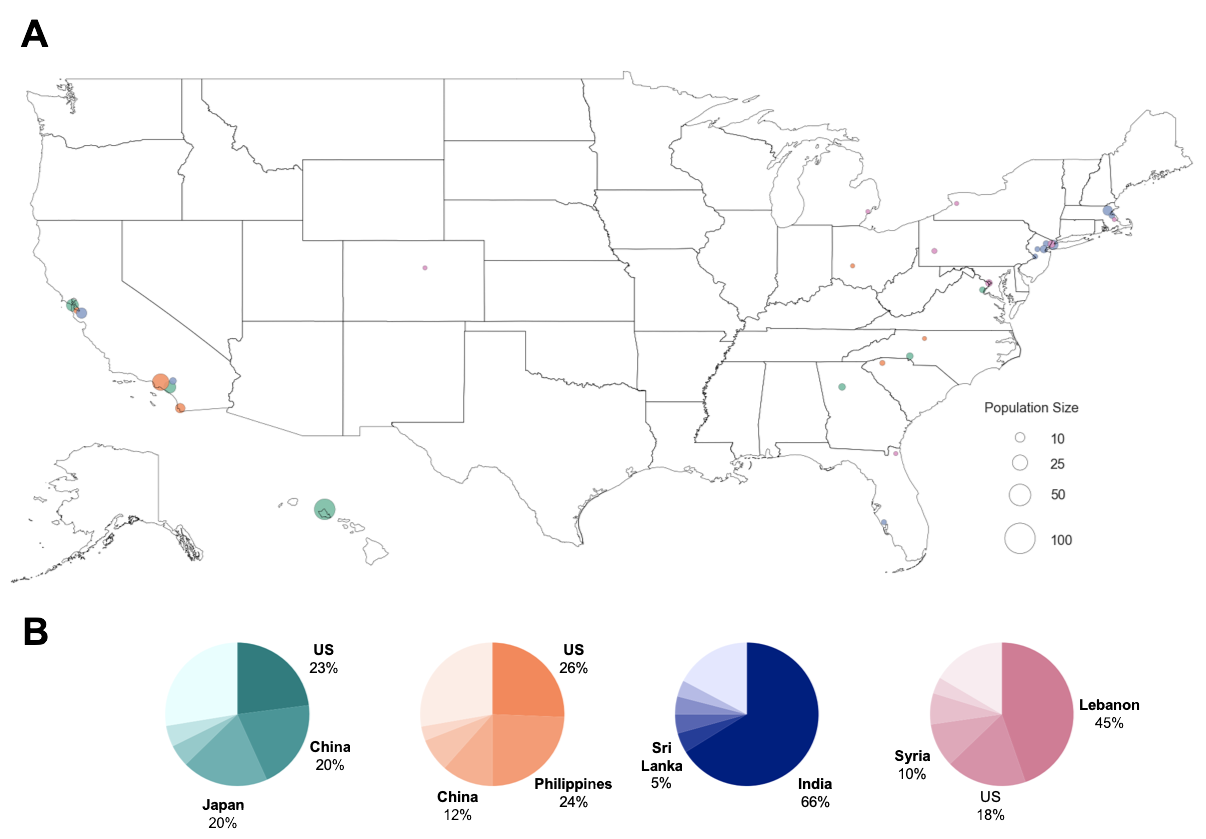

**Figure S10. Distribution of Haplotype Clusters with Asian Ancestries**

**(A)** Map of haplotype clusters corresponding to regional Asian ancestries. Each county containing present-day individuals is represented by a dot. The top 10 locations with the highest odds ratio are shown for each cluster. Maps showing the full distribution for each cluster can be found in the supplement (**Figure S11**).

**(B)** Ancestral birth origin proportions for each cluster in (A). Only individuals with complete pedigree annotations, up to grandparent level, are shown.

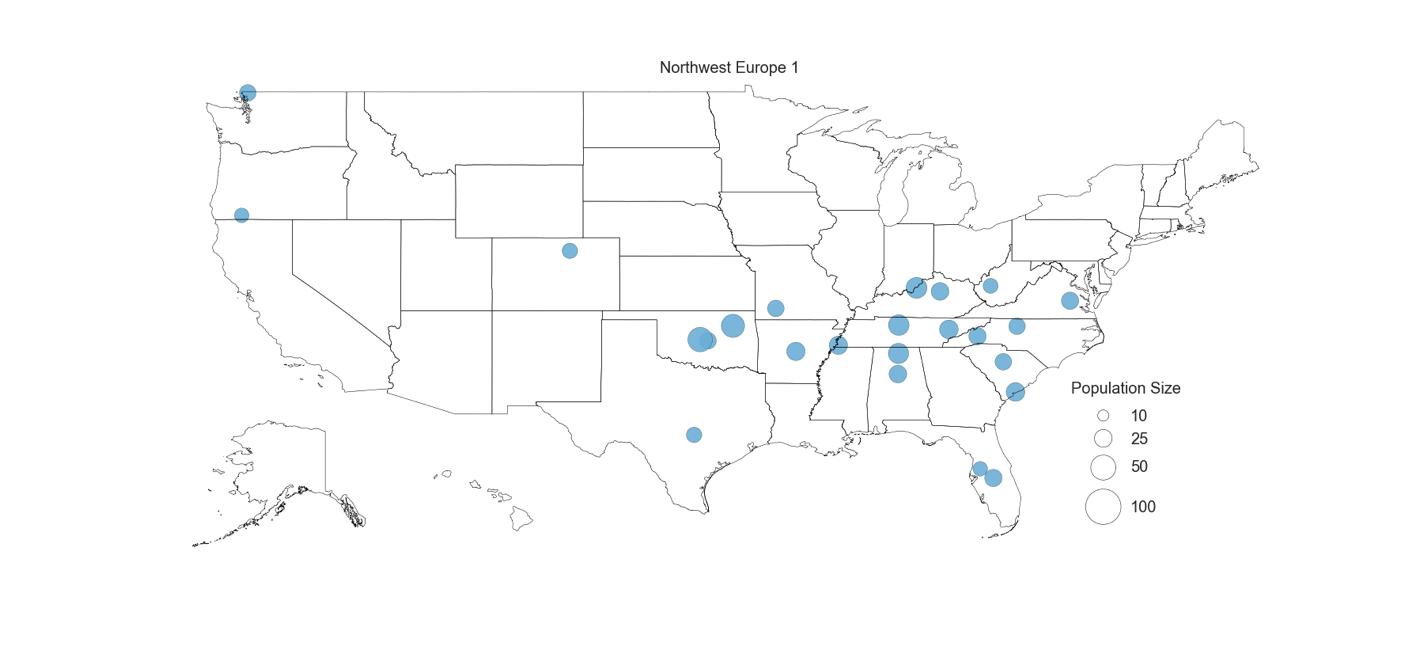

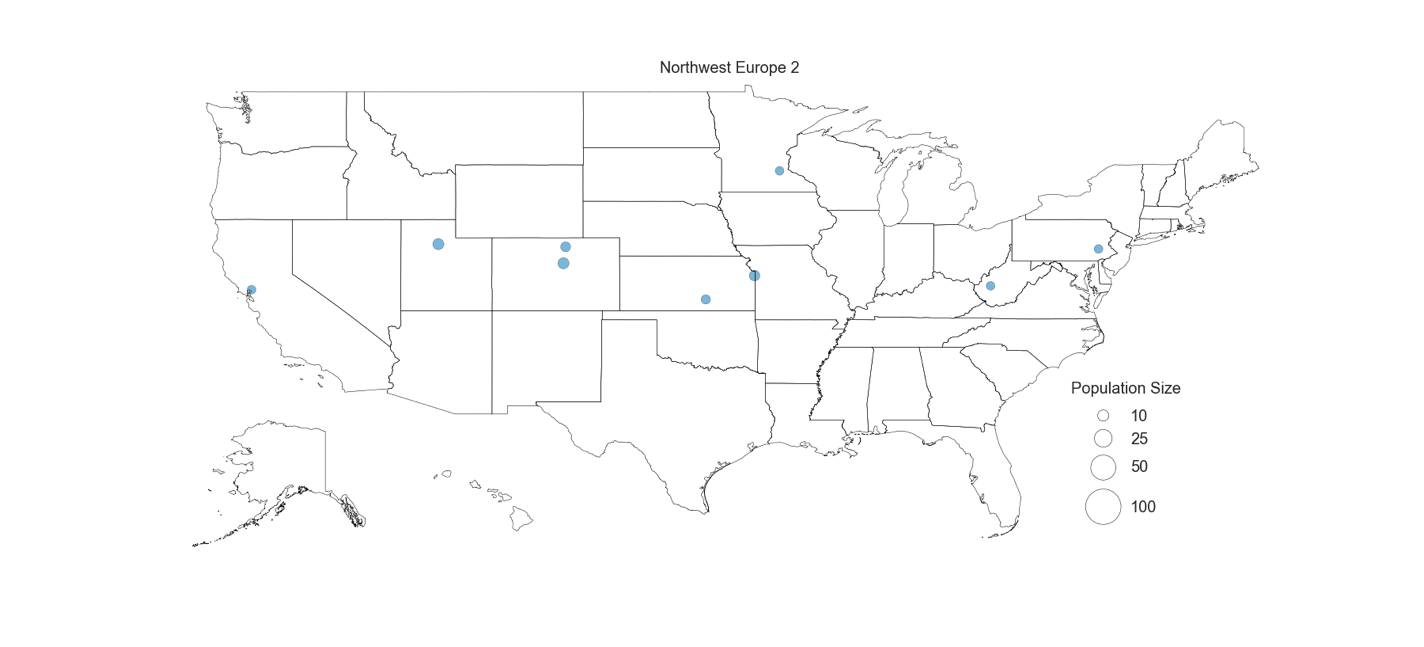

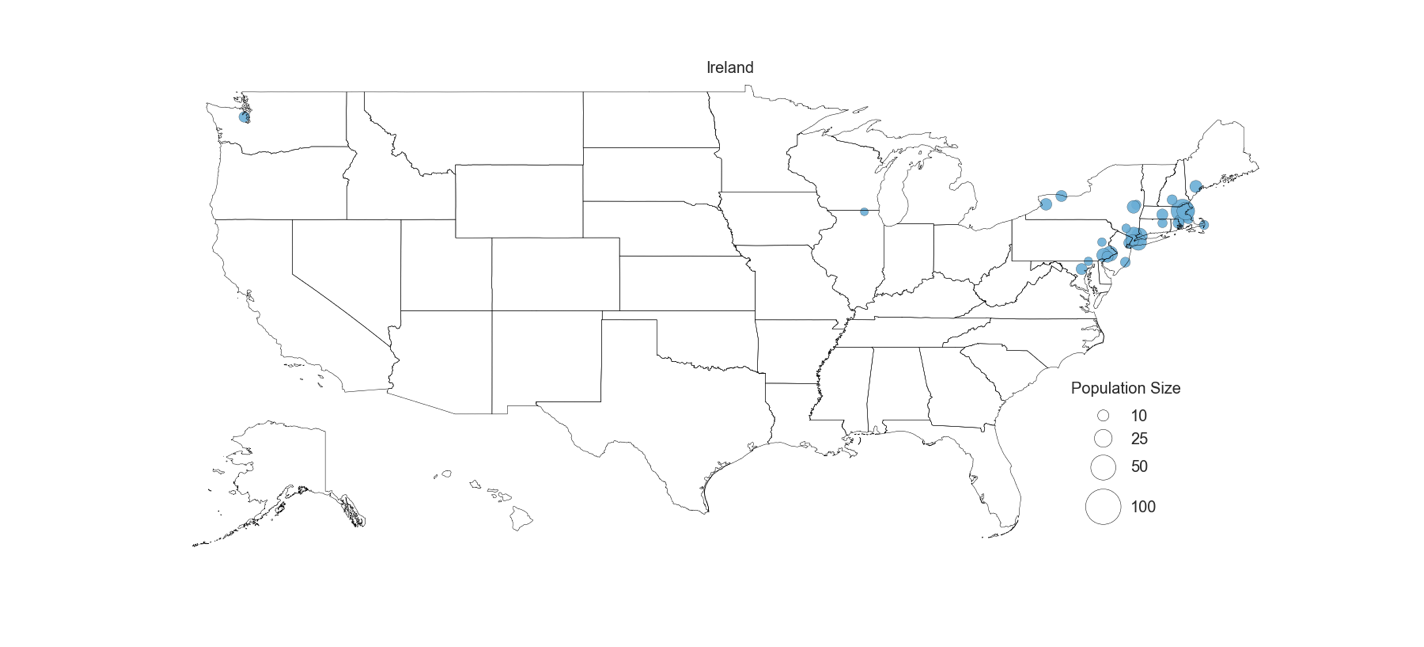

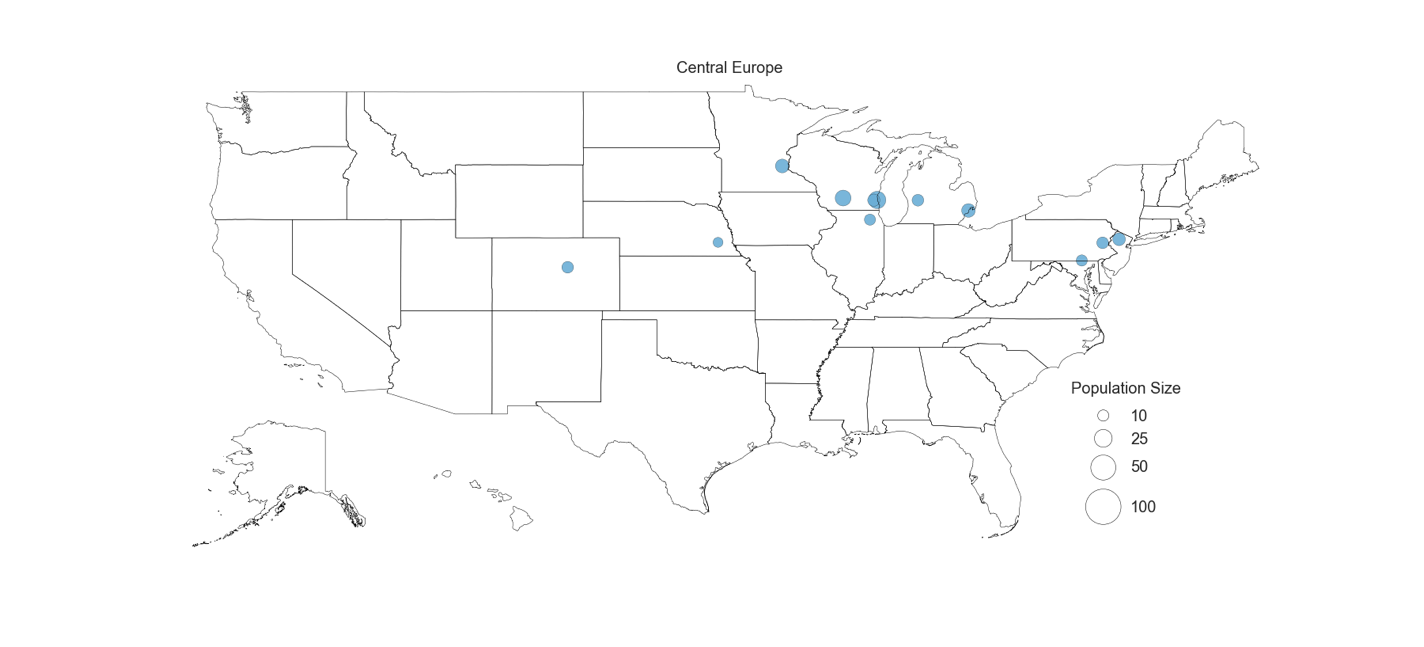

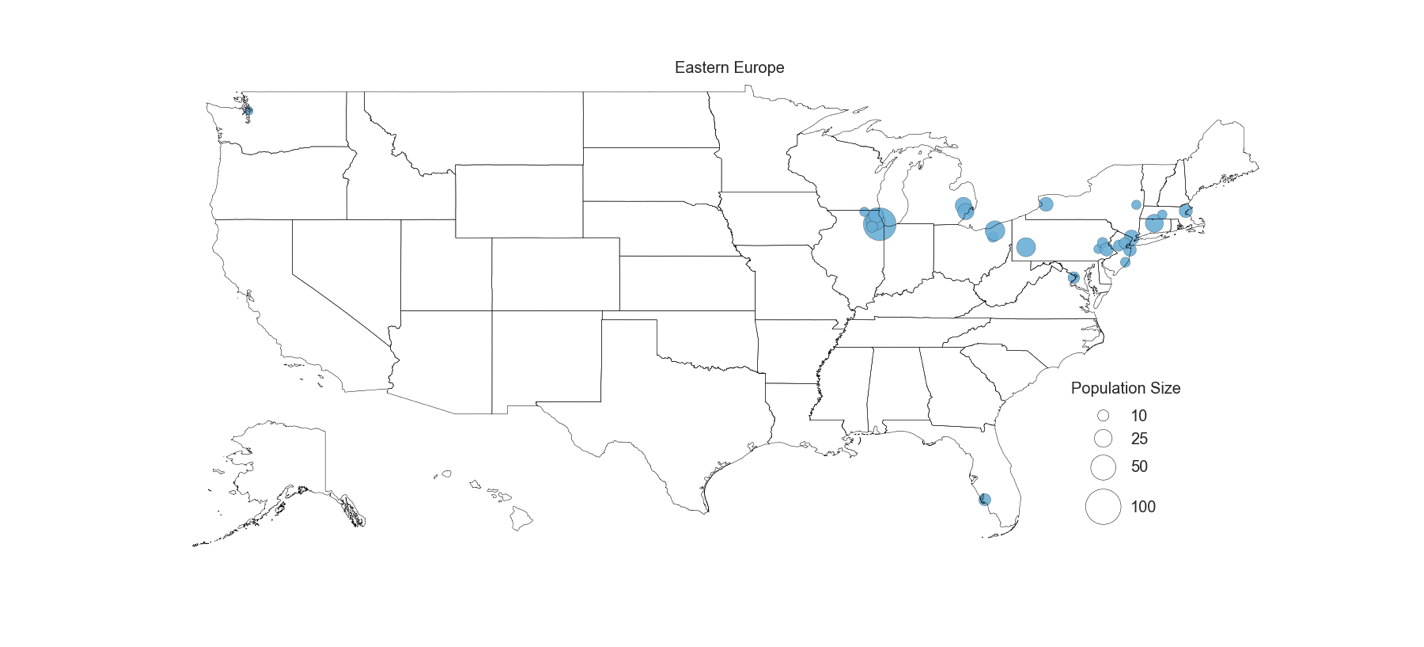

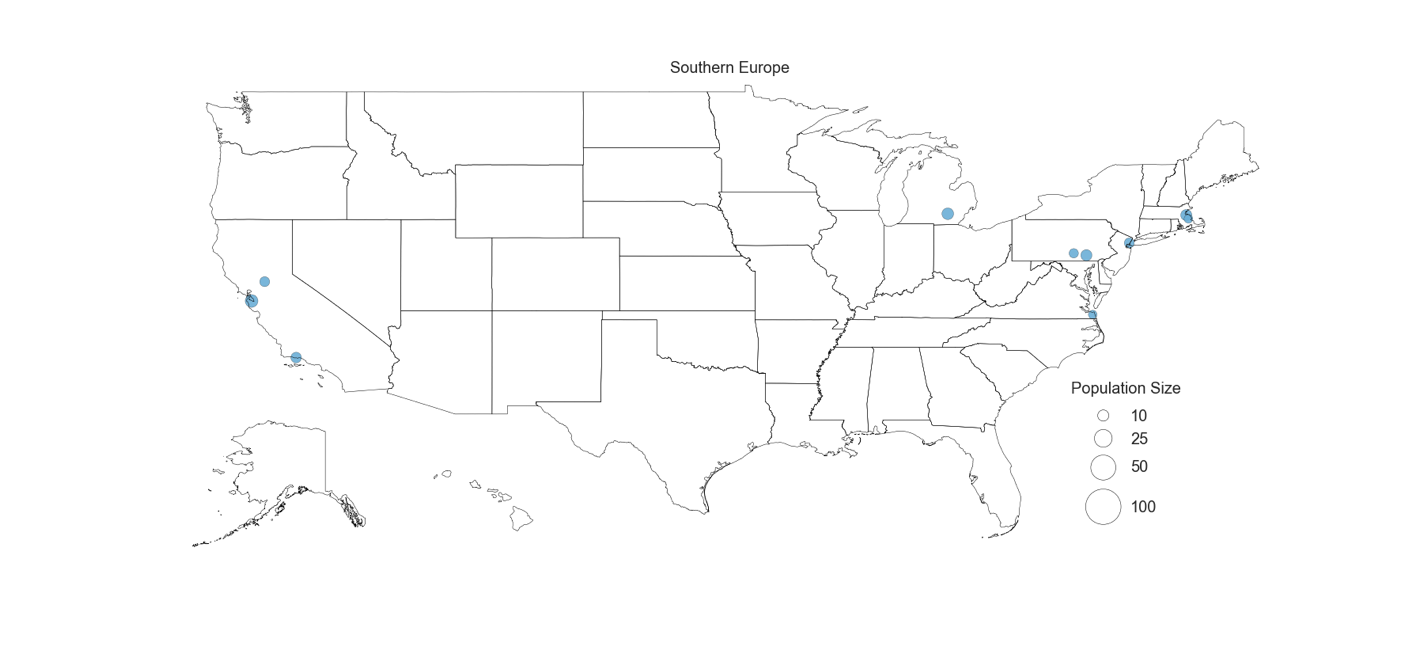

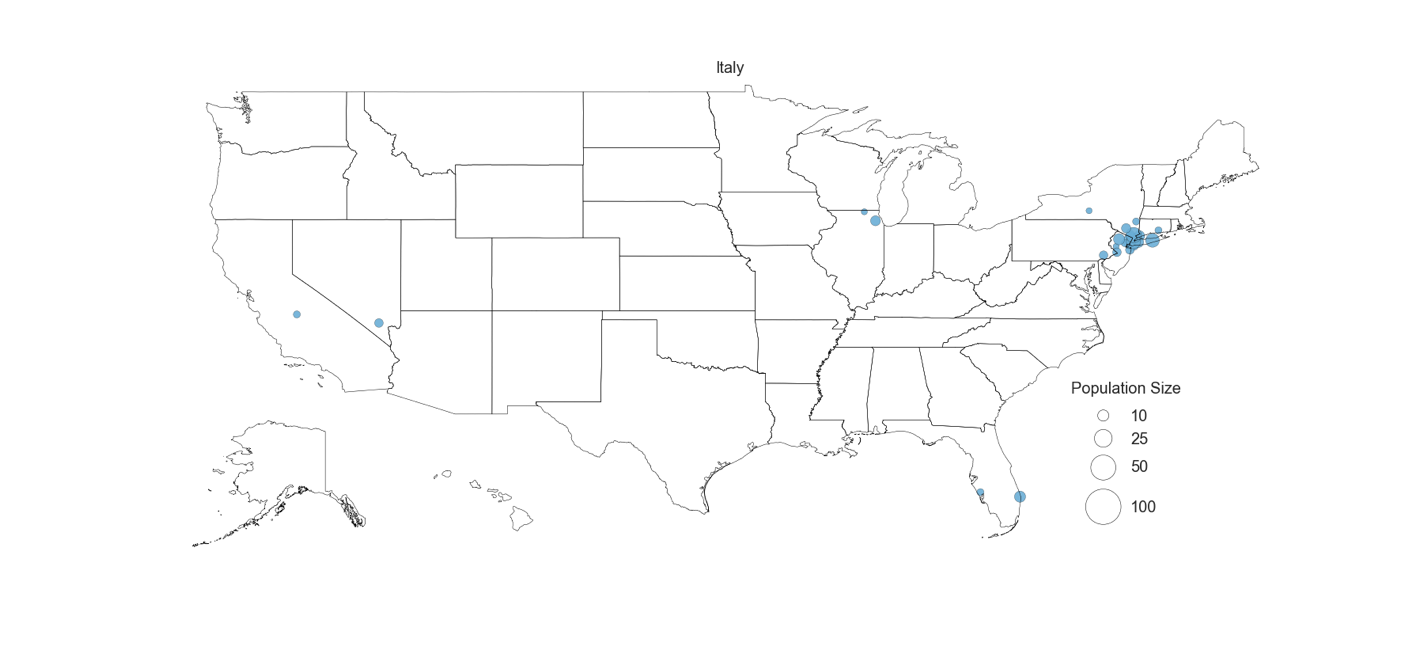

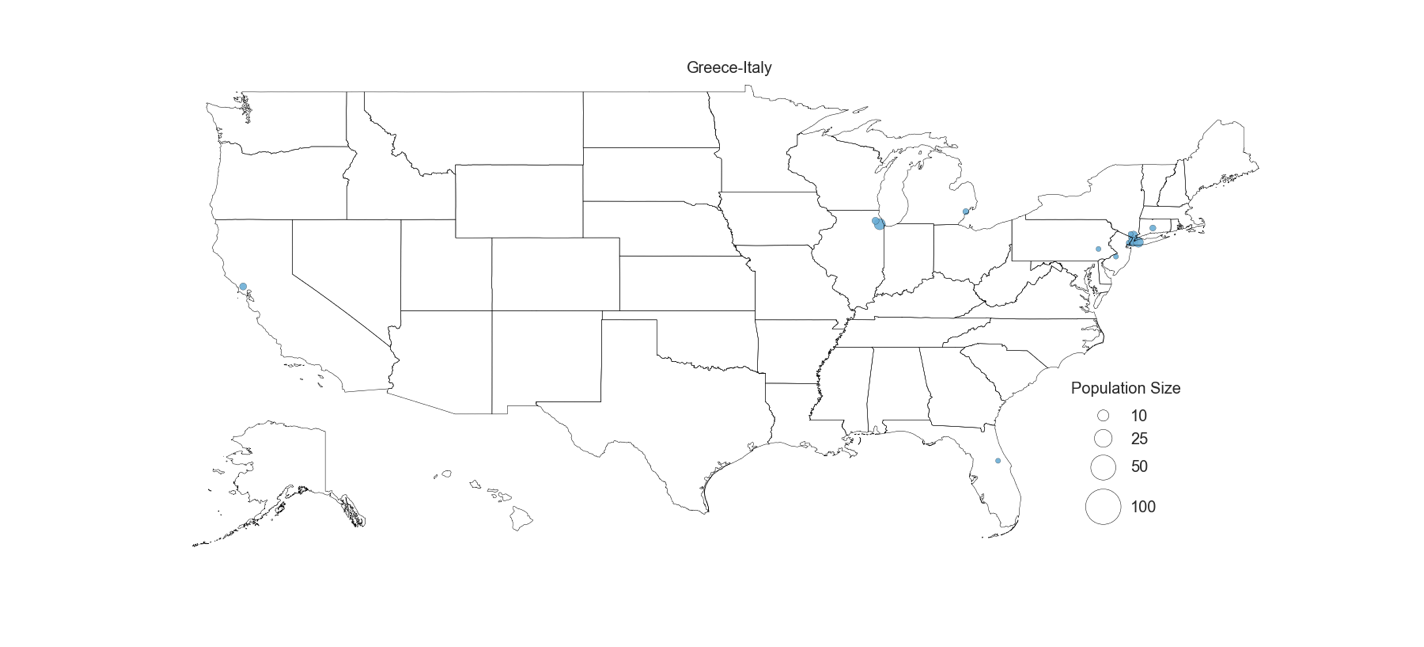

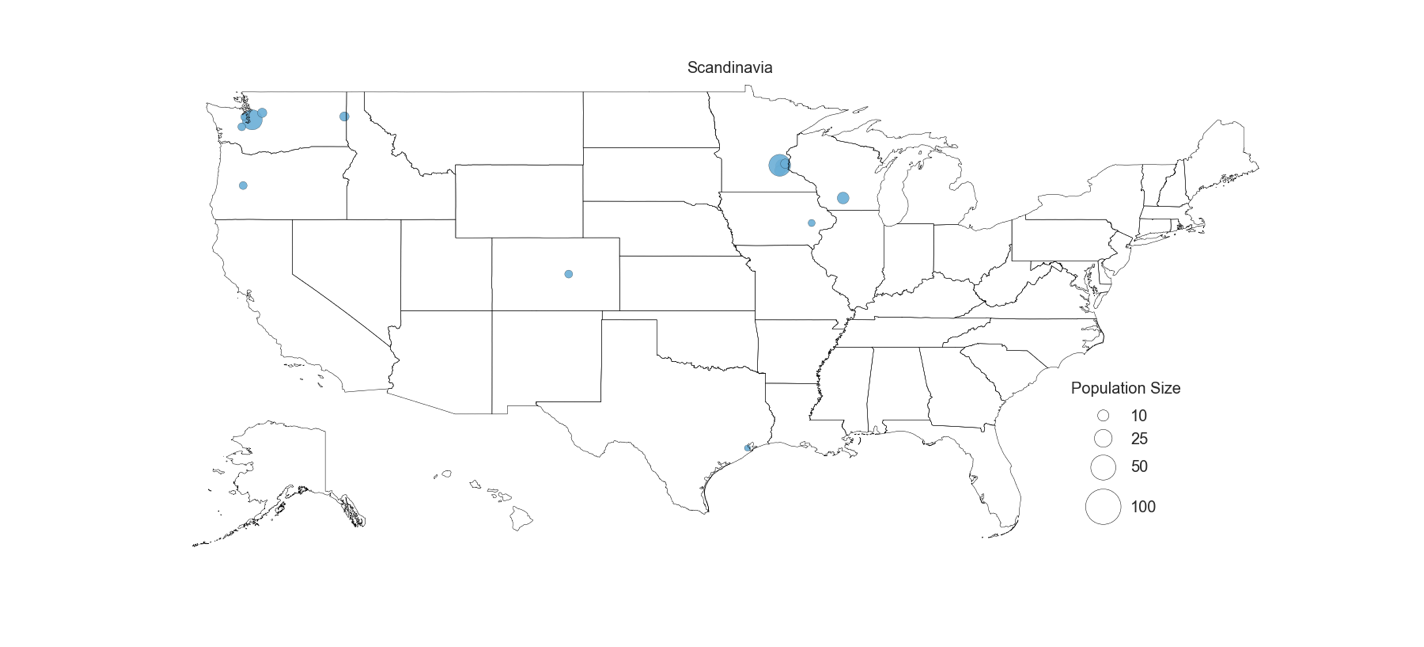

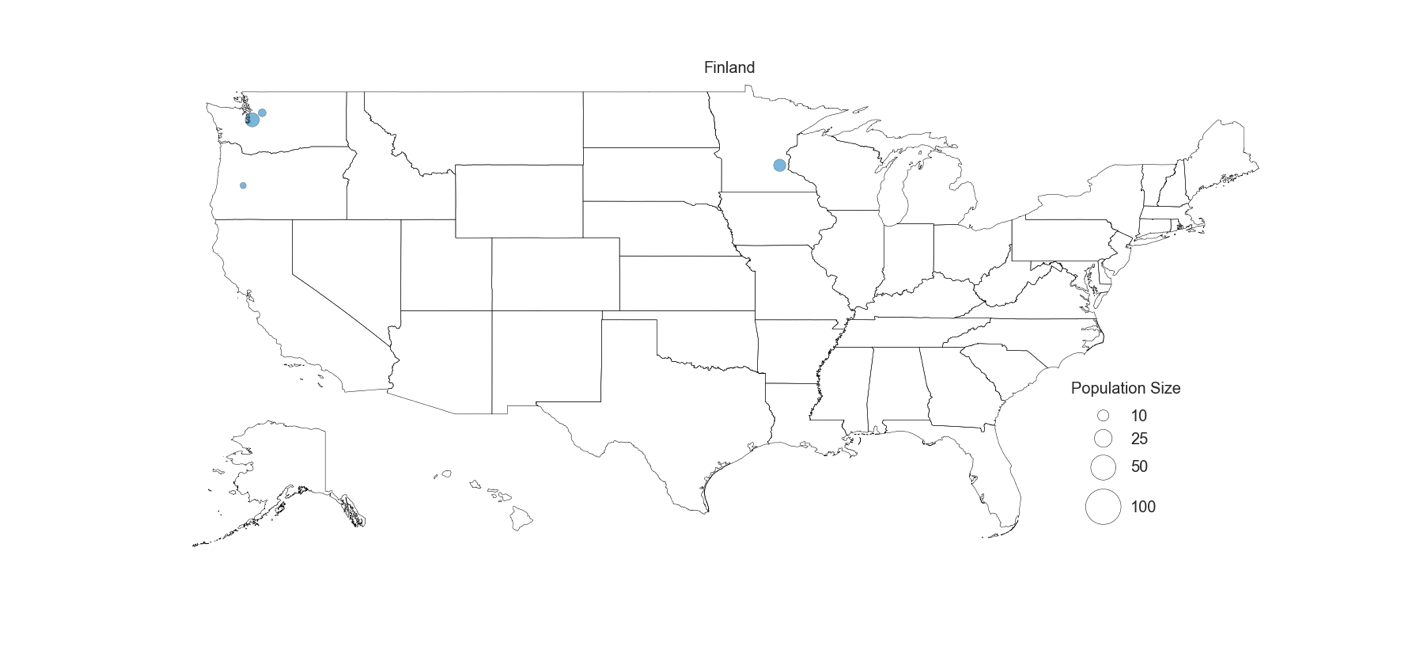

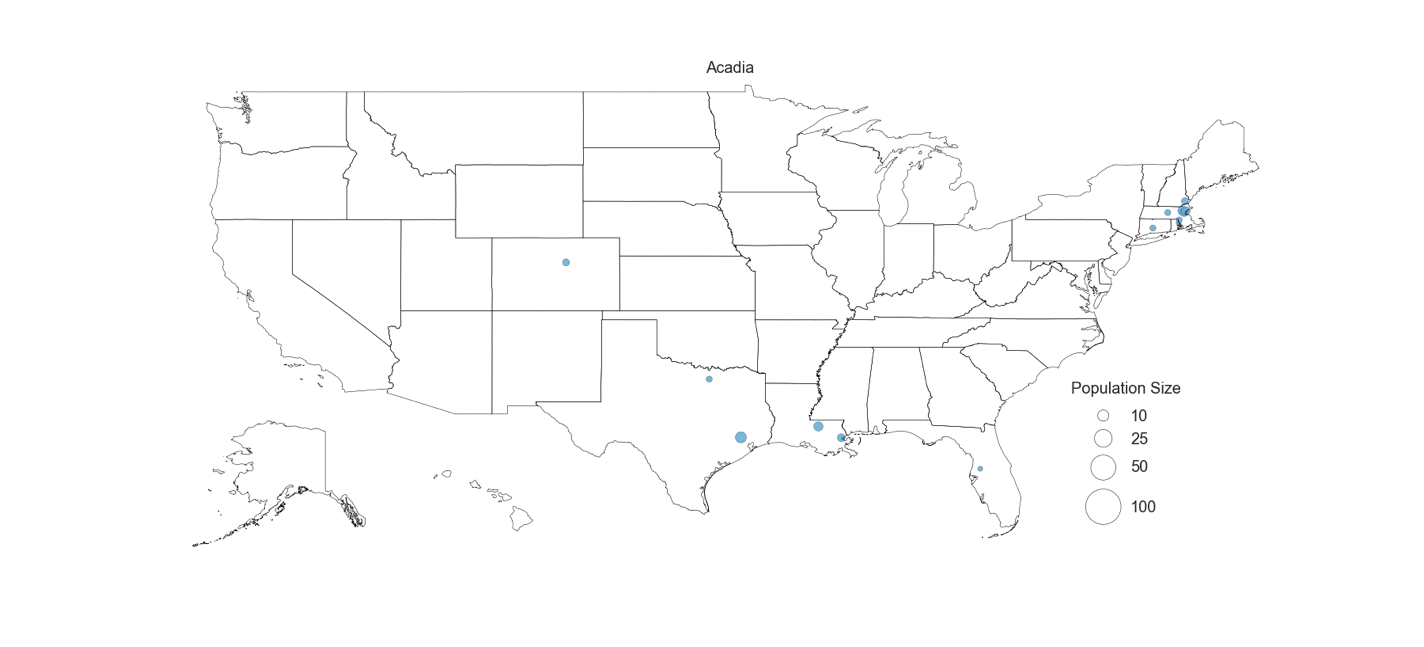

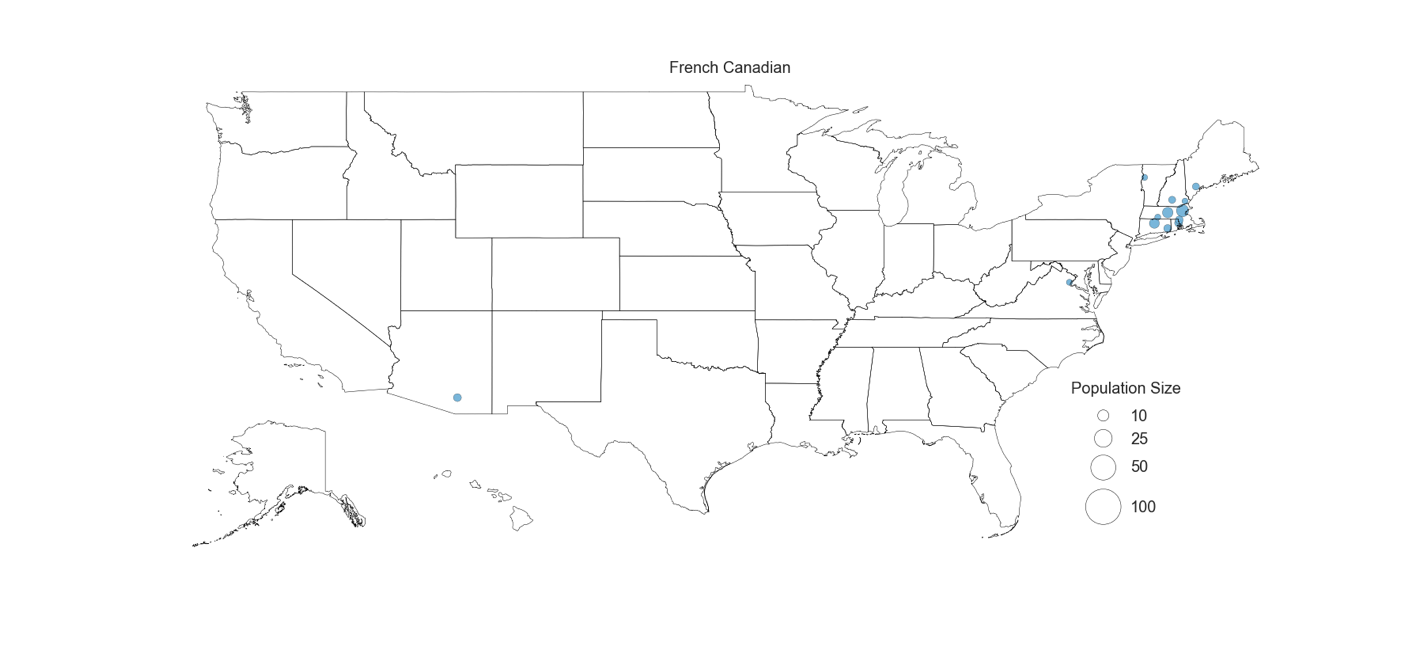

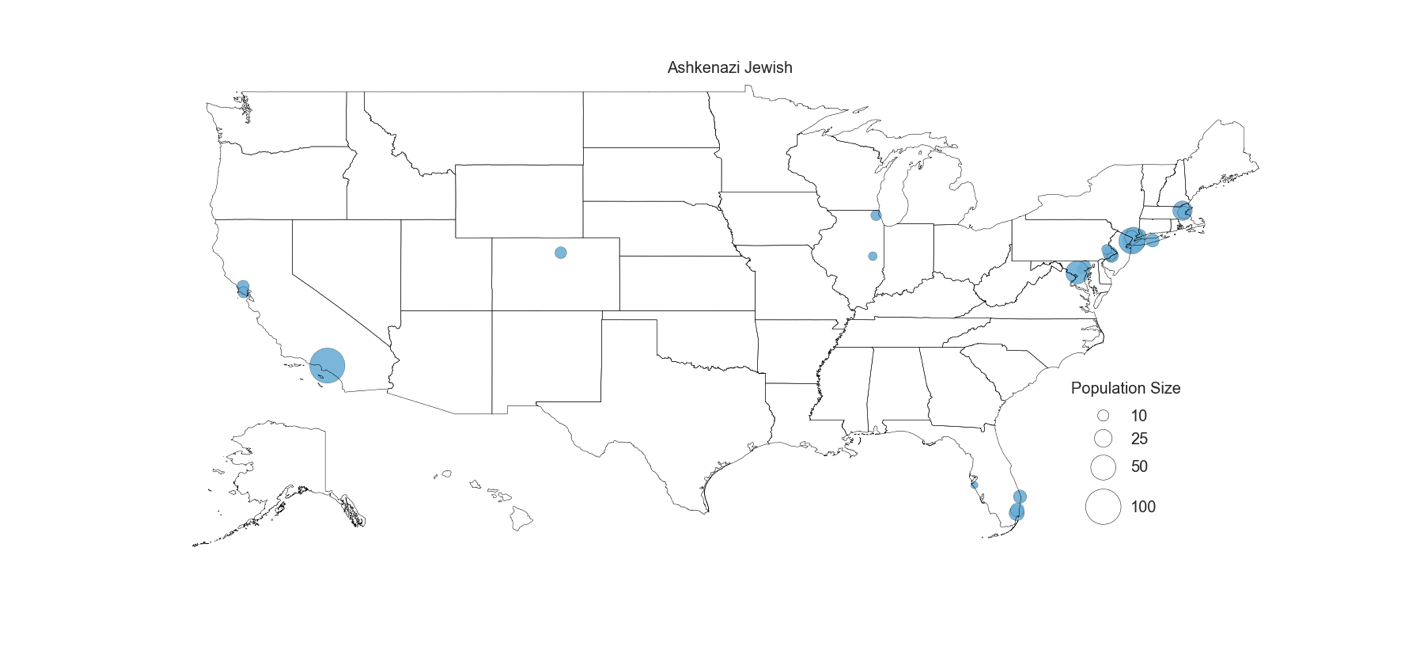

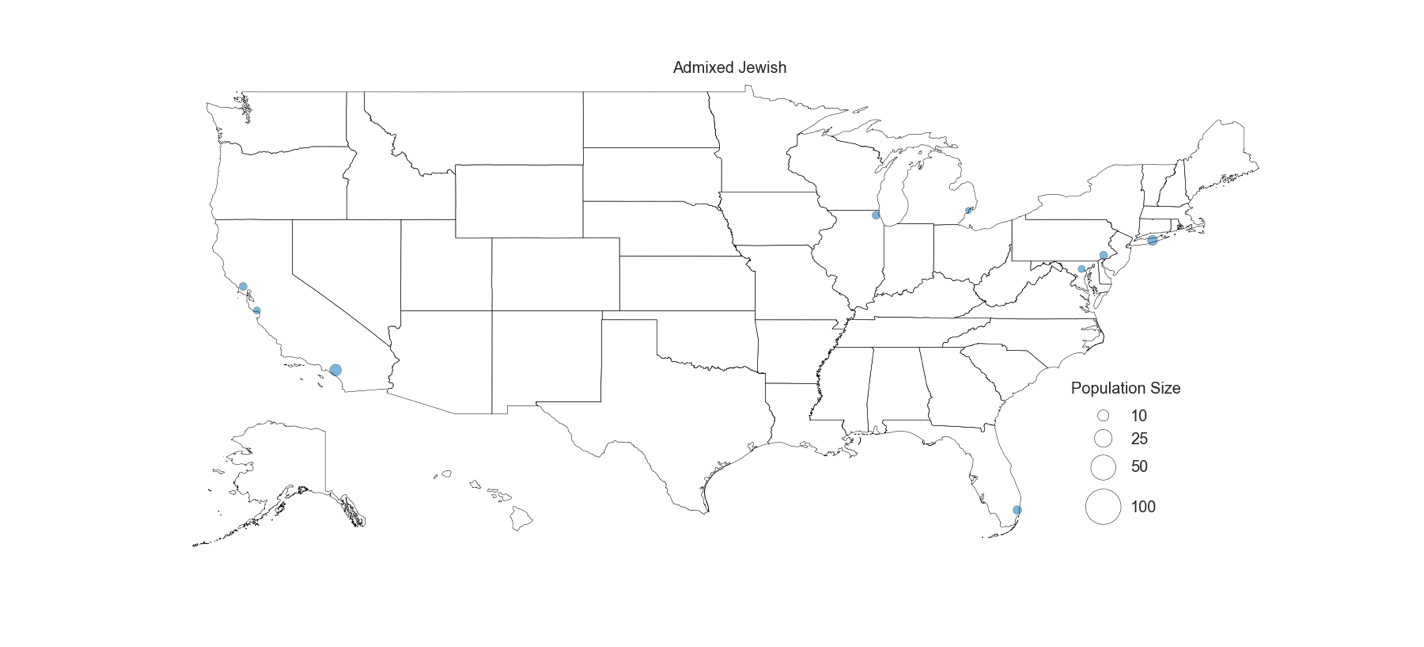

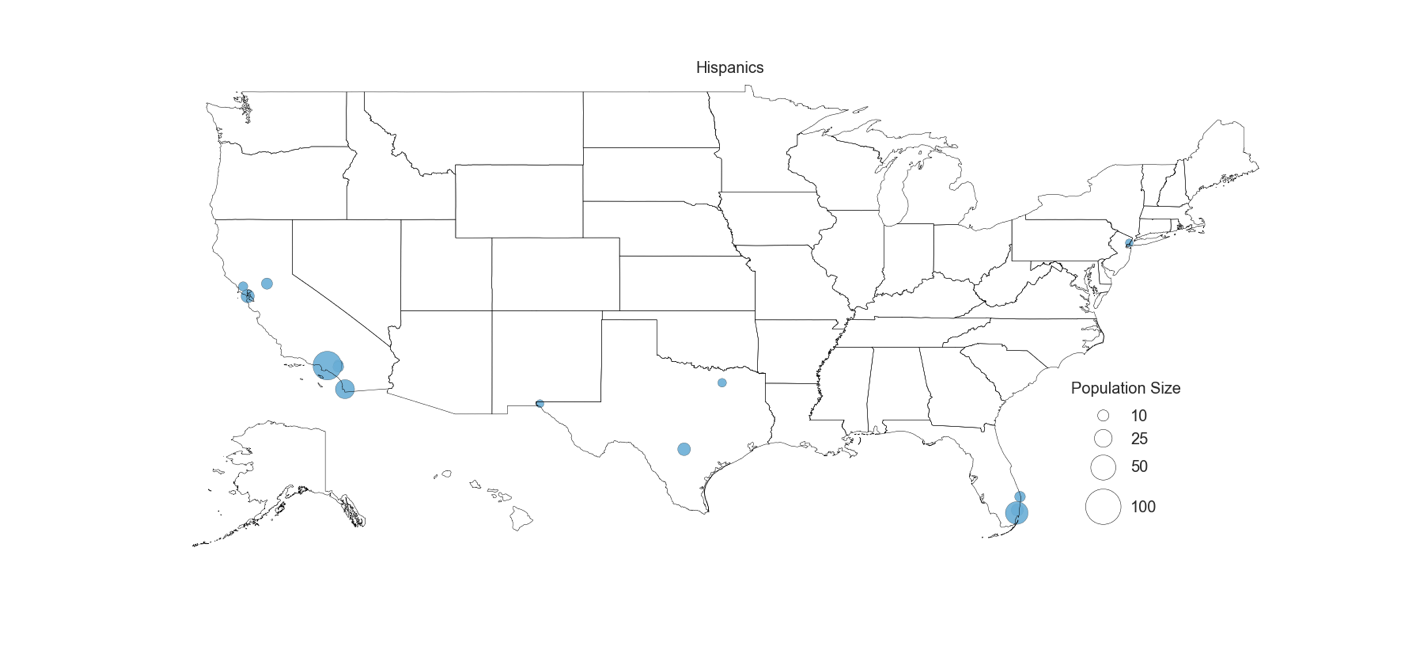

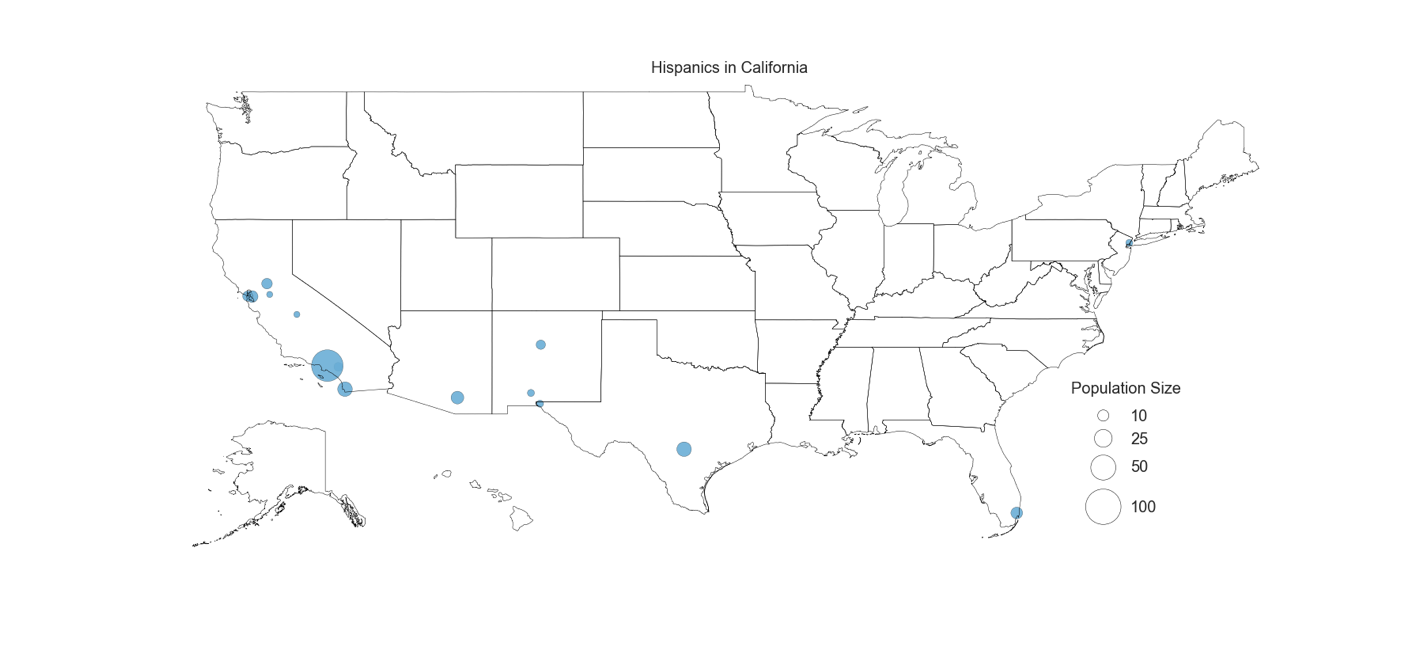

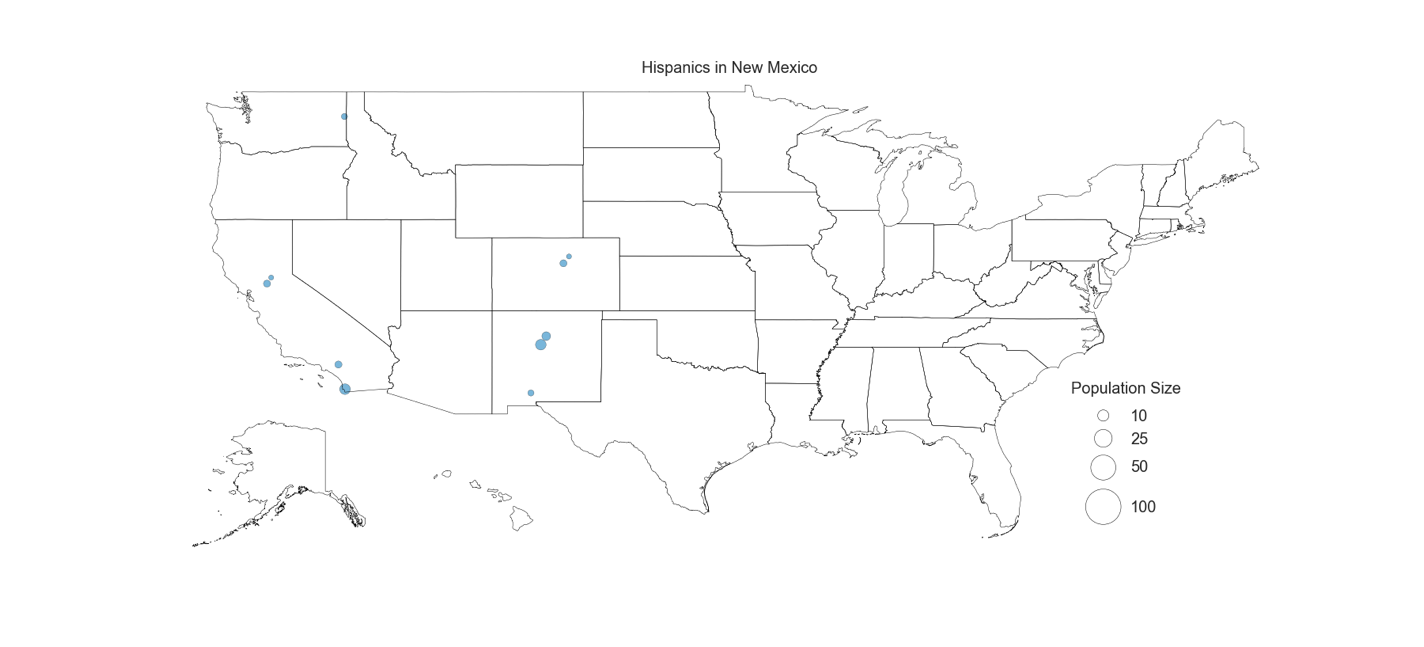

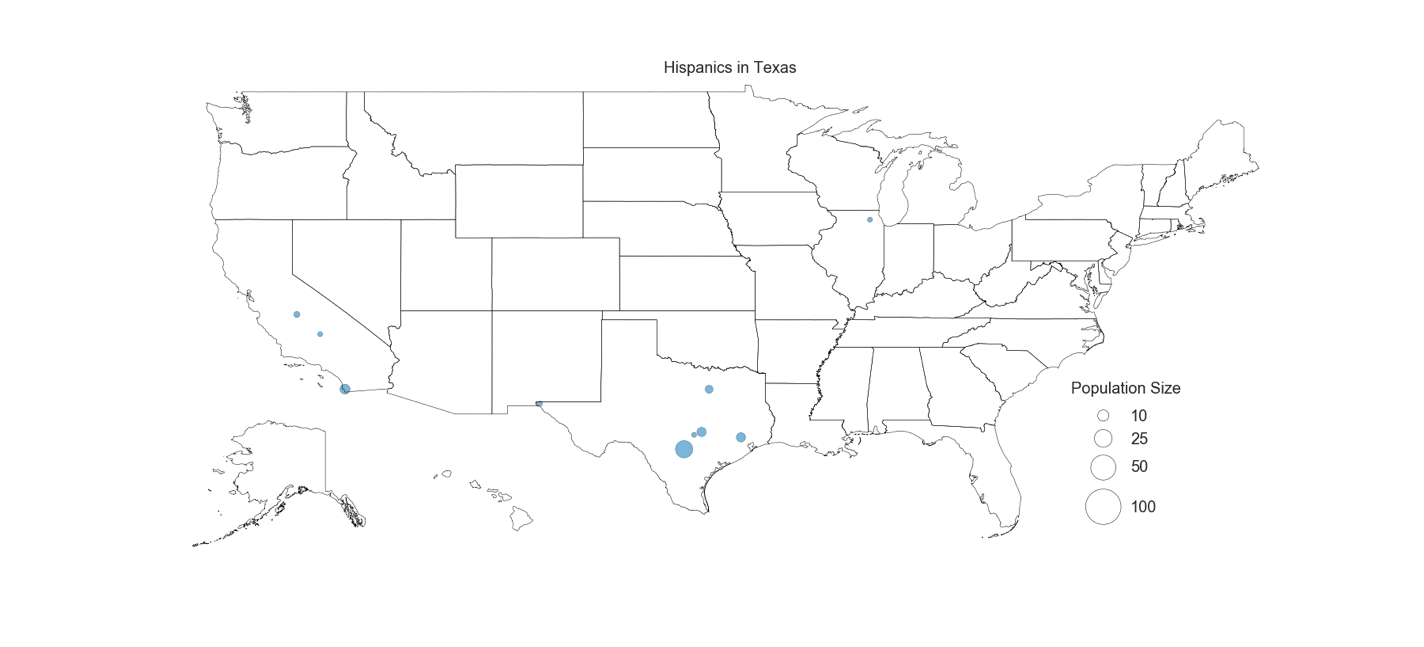

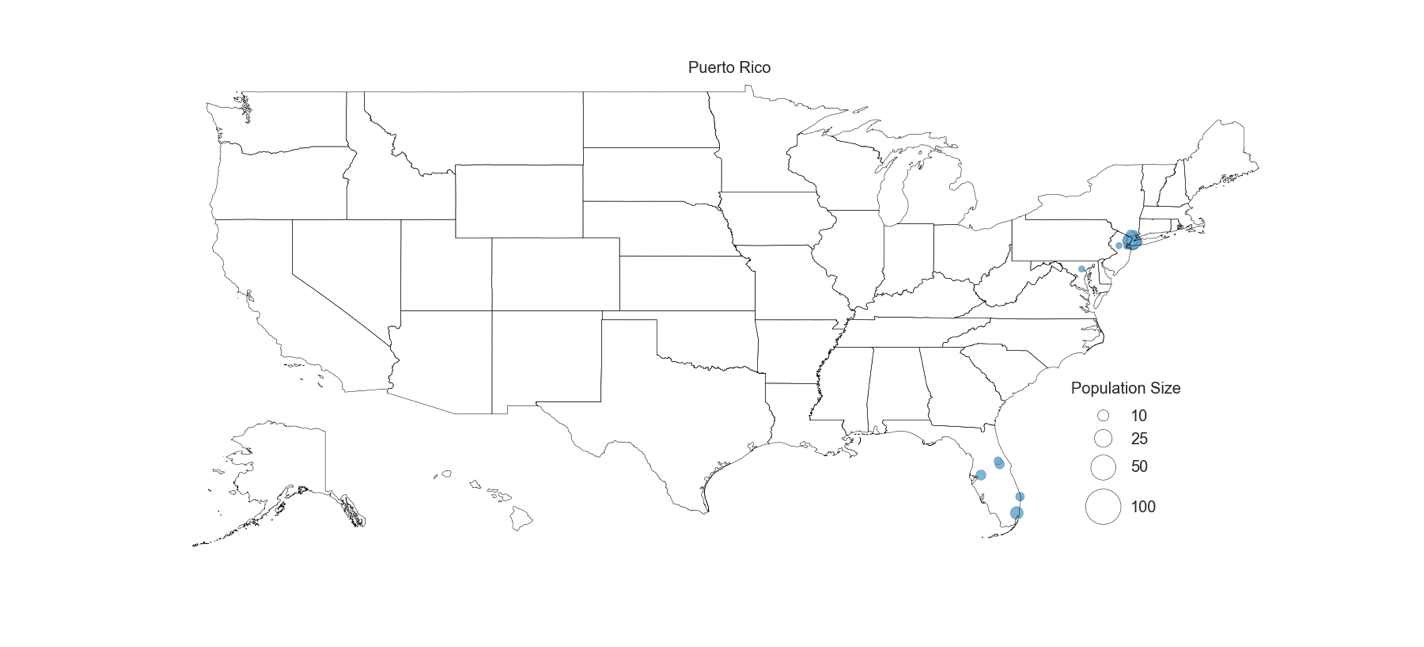

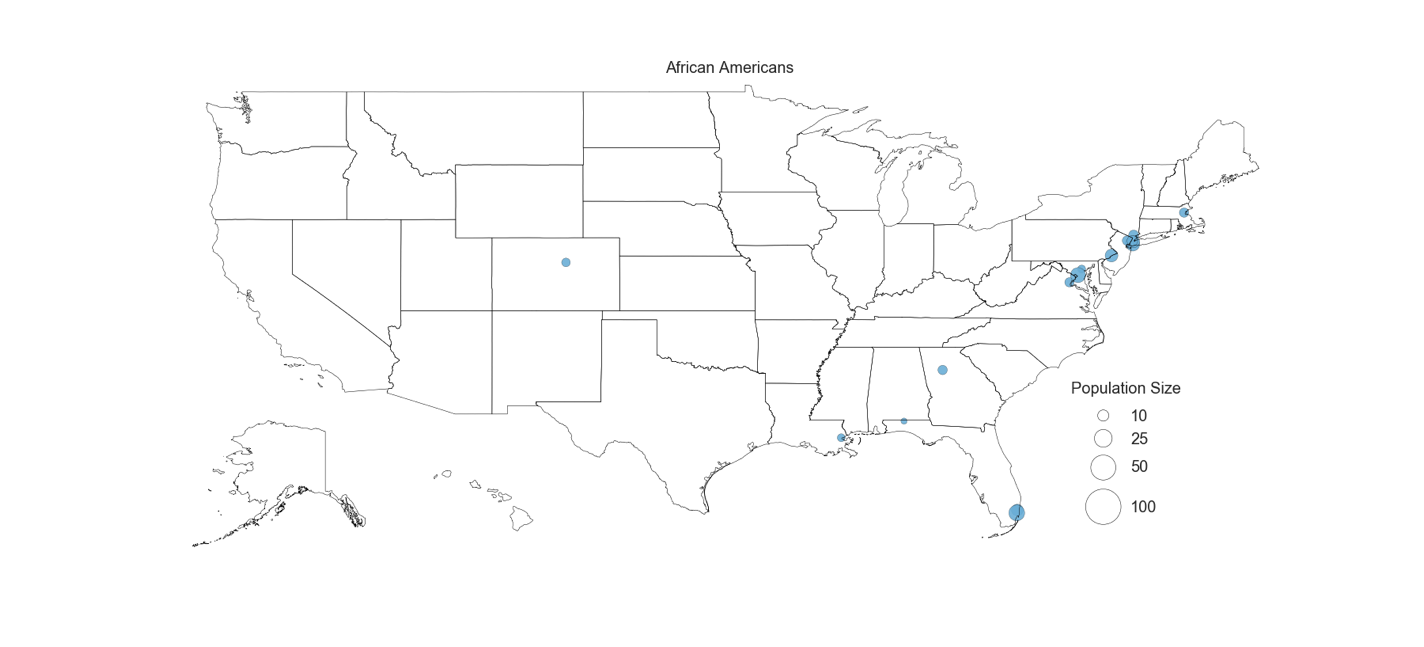

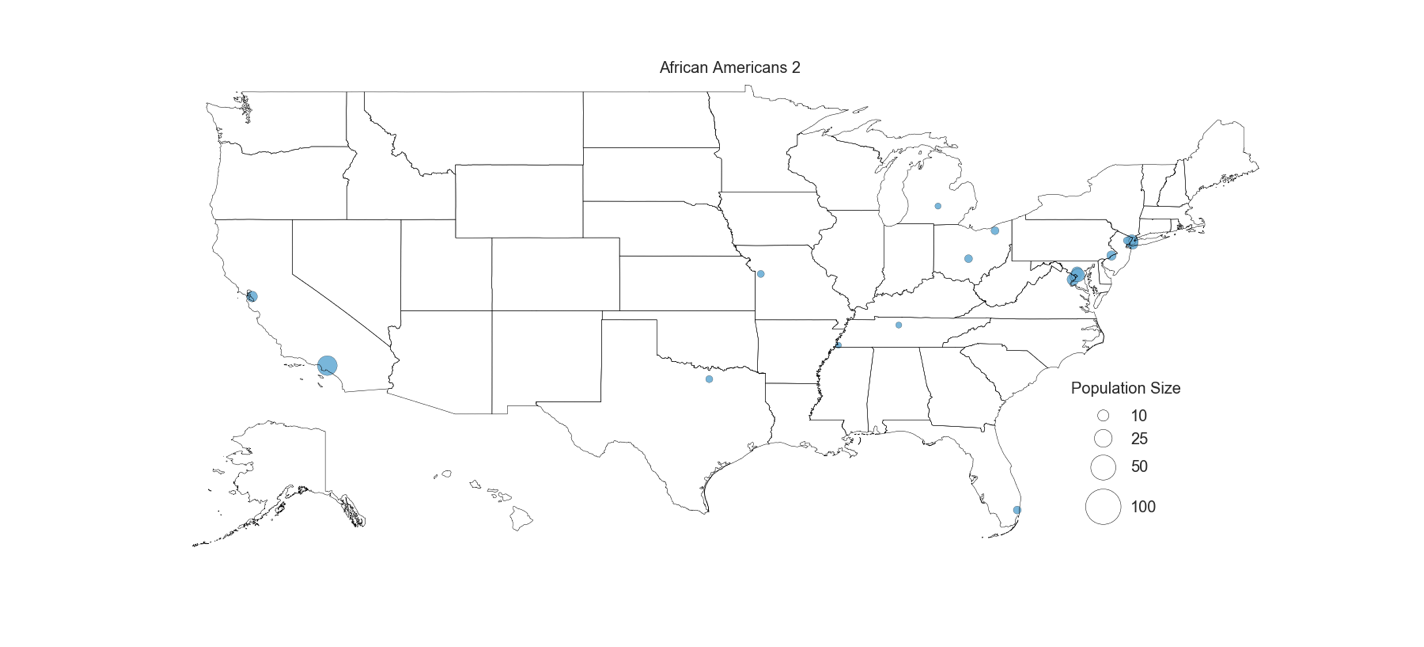

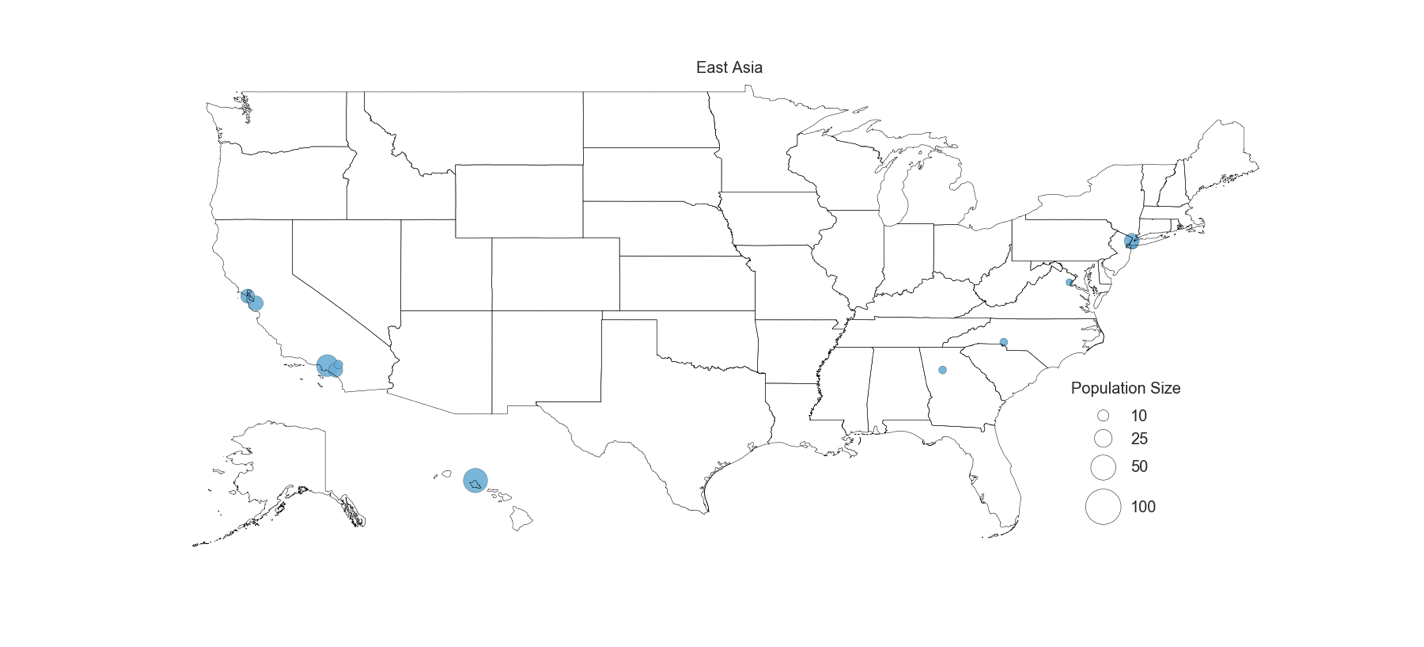

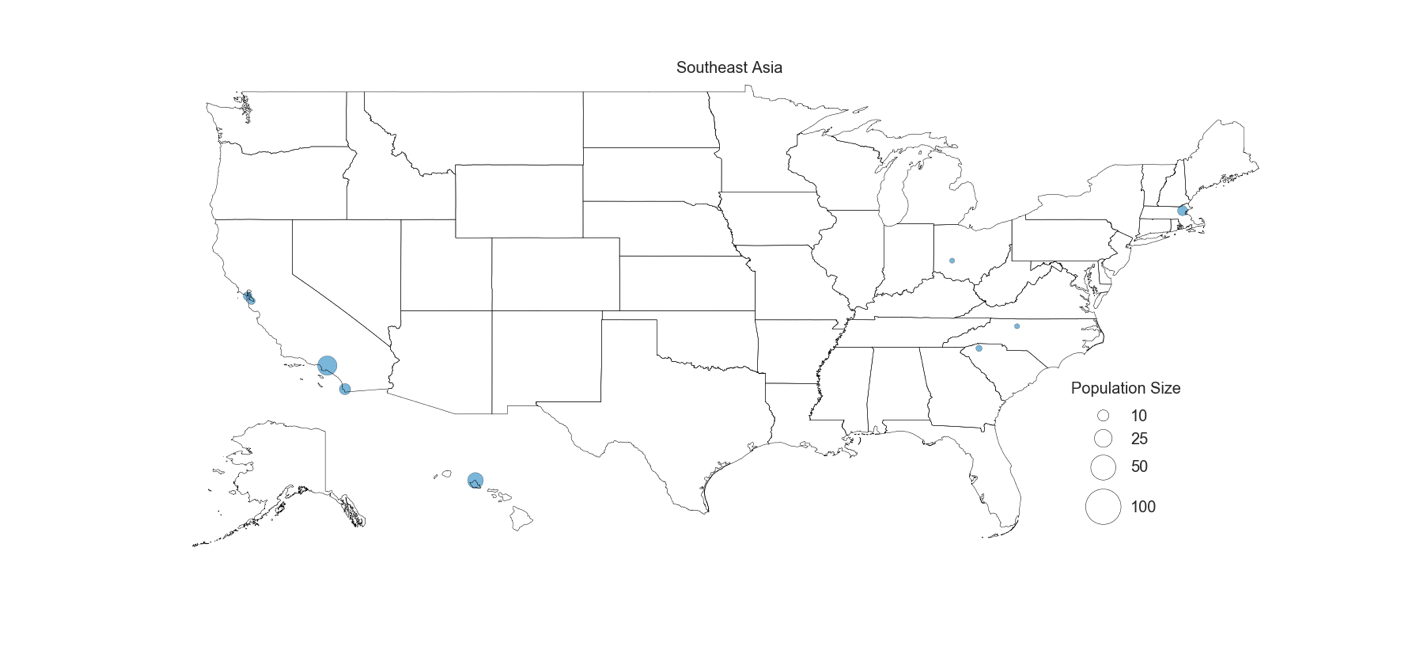

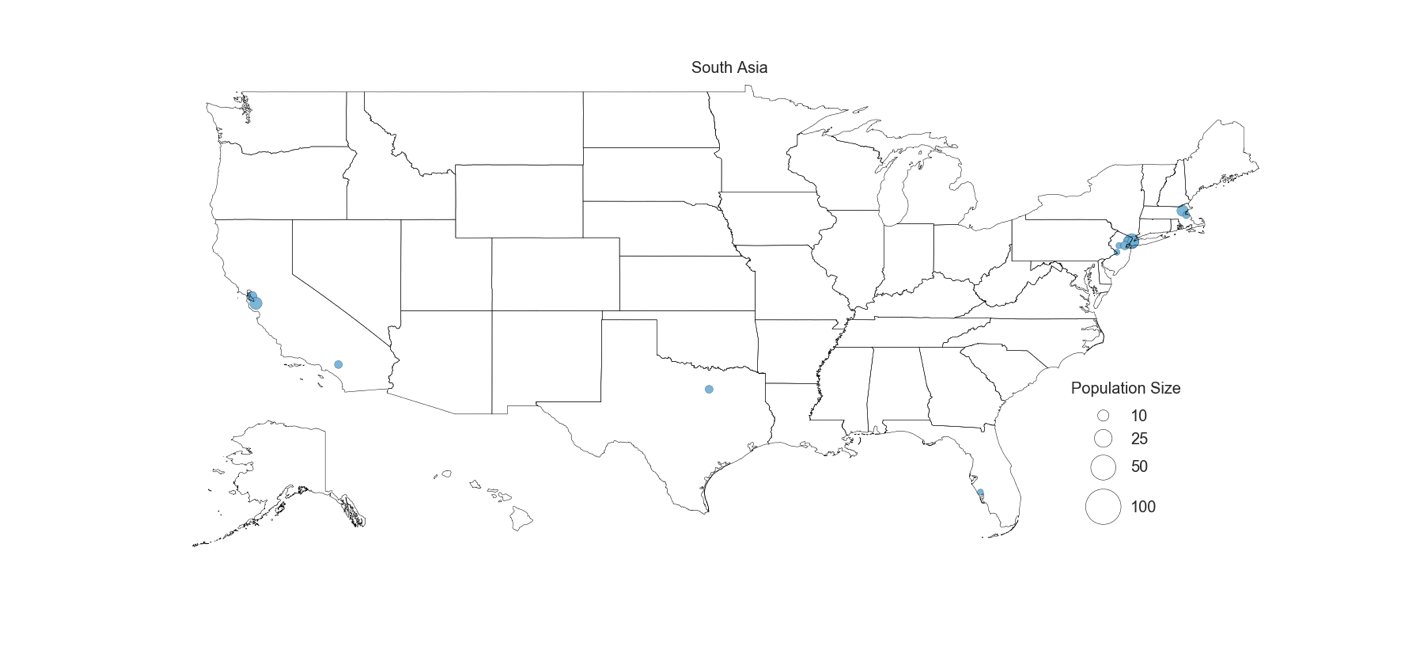

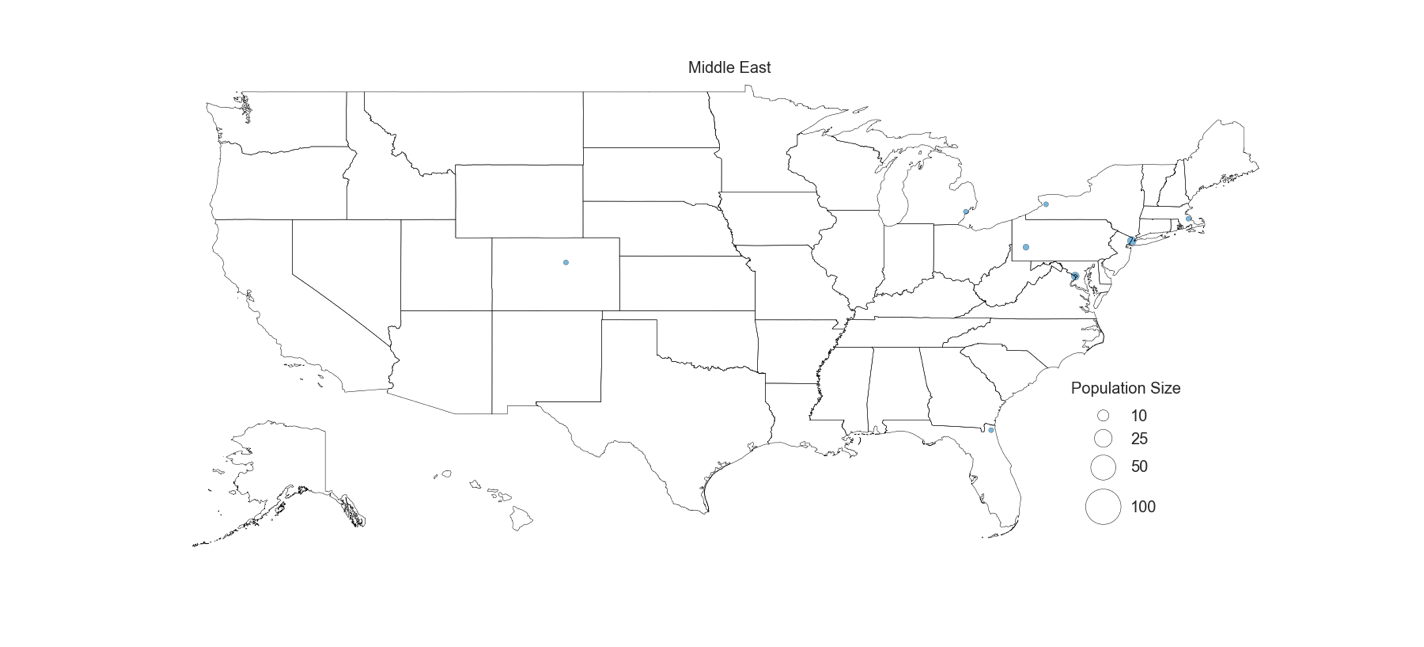

**Figure S11. Geographical Distribution of Haplotype Clusters**

Present-day location of individuals in each cluster. Each county is represented by a dot and only the cluster with the highest odds ratio. All counties with statistically significant presence of individuals are shown for each cluster.
